# The cellular modifier MOAG-4/SERF drives amyloid formation through charge complementation

**DOI:** 10.1101/2020.12.09.417709

**Authors:** Anita Pras, Bert Houben, Francesco A. Aprile, Renée Seinstra, Rodrigo Gallardo, Leen Janssen, Wytse Hogewerf, Matthias De Vleeschouwer, Alejandro Mata-Cabana, Mandy Koopman, Esther Stroo, Minke de Vries, Samantha Louise Edwards, Michele Vendruscolo, S. Fabio Falsone, Frederic Rousseau, Joost Schymkowitz, Ellen A. A. Nollen

## Abstract

While aggregation-prone proteins are known to accelerate ageing and cause age-related diseases, the cellular mechanisms that drive their cytotoxicity remain unresolved. The orthologous proteins MOAG-4, SERF1A and SERF2 have recently been identified as cellular modifiers of such cytotoxicity. Using a peptide array screening approach on human amyloidogenic proteins, we found that SERF2 interacted with specific patterns of negatively charged and hydrophobic, aromatic amino acids. The absence of such patterns, or the neutralization of the positive charge in SERF2, prevented these interactions and abolished the amyloid-promoting activity of SERF2. In a protein aggregation model in the nematode *C. elegans*, protein aggregation was suppressed by mutating the endogenous locus of MOAG-4 to neutralize charge. Our data indicate that charge interactions are required for MOAG-4 and SERF2 to promote aggregation. Such charged interactions might accelerate the primary nucleation of amyloid by initiating structural changes and by decreasing colloidal stability. Our finding that negatively charged segments are overrepresented in amyloid-forming proteins suggests that inhibition of charge interactions deserves exploration as a strategy to target age-related protein toxicity.

**Significance Statement:** How aging causes relatively common diseases such as Alzheimer’s and Parkinson’s is still a mystery. Since toxic structural changes in proteins are likely to be responsible, we investigated biological mechanisms that could drive such changes. We made use of a modifying factor called SERF2, which accelerates structural changes and aggregation of several disease-related proteins. Through a peptide-binding screen, we found that SERF2 acts on negatively charged protein regions. The abundance of such regions in the disease-related proteins explains why SERF has its effect. Removing positive charge in SERF was sufficient to suppress protein aggregation in models for disease. We propose that blocking charge-interactions with SERF or other modifiers could serve as a general approach to treat age-related protein toxicity.

## Introduction

Protein homeostasis declines with ageing (1–4). This decline results in an increased accumulation of aggregation-prone proteins, which accelerates ageing and is associated with a wide range of age-related disorders, including Alzheimer’s and Parkinson’s diseases. Although the different proteins involved in these diseases are usually unrelated in sequence and native structure, they share the tendency to convert into ordered, cross-ß structures known as amyloid fibrils (5–7). Structural conversions early in the aggregation process play an important role in amyloid formation and the associated cellular toxicity (8–10). The cellular mechanisms that drive these early structural conversions, however, are poorly understood, and uncovering them is key for the development of interventions to prevent such toxic structural changes in amyloid-associated diseases.

For several structurally unrelated amyloidogenic proteins, previous studies have identified proteins that enhance such structural conversions, namely modifier of aggregation-4 (MOAG-4) in the nematode worm *Caenorhabditis elegans* and its human orthologs small EDRK-rich factors (SERF) 1A and 2 (11–13). *In vitro* studies with purified proteins have shown that SERF1A preferentially promotes the aggregation of amyloidogenic proteins – including alpha-synuclein, amyloid beta and prion protein – above the aggregation of non-amyloidogenic proteins (13). MOAG-4/SERF1A is involved early on in the aggregation process. In the case of alpha-synuclein, MOAG-4 and SERF hinder intermolecular interactions within the protein, through electrostatic interactions (11, 14). This results in a more aggregation-prone conformation of alpha-synuclein, which in turn seeds the formation of amyloid fibrils that can eventually form large, insoluble aggregates (11, 13, 14). In addition, previous studies have shown that MOAG-4 and SERF1A act transiently and are not incorporated in amyloid fibrils themselves (11, 13). The shared mechanism by which MOAG-4 and SERFs bind to and induce the structural conversion of other amyloidogenic proteins remains nevertheless unknown.

We therefore use a peptide-array screening approach to identify interactions that accelerate the aggregation of amyloidogenic proteins. Our findings suggest that SERF2 affects the earliest steps of the aggregation process by charge complementation on amyloidogenic proteins, thereby inducing structural conversions that accelerate fibril formation.

## Results

### SERF2 selectively binds to negatively charged peptides

To determine how SERF catalyzes amyloid formation of multiple unrelated amyloidogenic proteins, we first used a peptide microarray-based approach to screen for SERF2-interacting amino acid sequences. The microarray contained peptide fragments from 27 full-length parent proteins and four dipeptide repeat polymers. Of these proteins, 19 are amyloidogenic (15) (Table S1, column A). Four of these amyloidogenic proteins have previously been shown to functionally interact with SERF (Table S1, proteins marked in bold) (12–14, 16). The other proteins represented on the slide included three other disease-related aggregation-prone proteins, four disease-related dipeptide repeat polymers, a protein for which amyloid formation is part of its physiological function and four non-amyloidogenic proteins (Table S1, columns B, C, D).

The full-length amino acid sequences for each of these proteins were represented by 12-mer peptides that overlapped by eight residues, hence producing a sliding window over each protein sequence (Figure 1A). The microarray contained duplicates of each peptide, randomly distributed over the array (Figure 1A and Dataset S1). Binding of ATTO633-labeled SERF2 (Uniprot identifier P84101-1, 59 amino acids) was visualized by fluorescent laser scanning and quantified based on fluorescence intensity (Dataset S1). Peptides were classified as SERF2 binders or non-binders based on their fluorescence intensity relative to a set of glycine controls (Gly) (Figure 1B and Dataset 1). Therefore, in each of three repeat experiments, the distribution of fluorescence intensities of the Gly control peptides was assessed (Figure 1B, upper panels). For each experiment, the cutoff was determined as the mean fluorescence of the Gly control peptides plus twice their standard deviation (red dashed lines in Figure 1B). In each separate repeat, peptides were then classified as binders when the RFU (Relative Fluorescence Units) values of both duplicates on the array were higher than the cutoff value sequences (Figure 1B and Dataset S1). Eventually, only peptides for which this was the case in all three experiments were classified as actual binders (Dataset S2). Peptides with RFUs below the threshold in each experiment were defined as non-binders, and peptides for which the classification varies between repeats were classified as “ambiguous”. In this way, 653 peptides were identified as binders, 2333 as non-binders, and 291 as ambiguous. These results show that SERF2 was able to bind selectively and reproducibly to a subset of the peptide sequences (Dataset S2).

**Figure 1.**
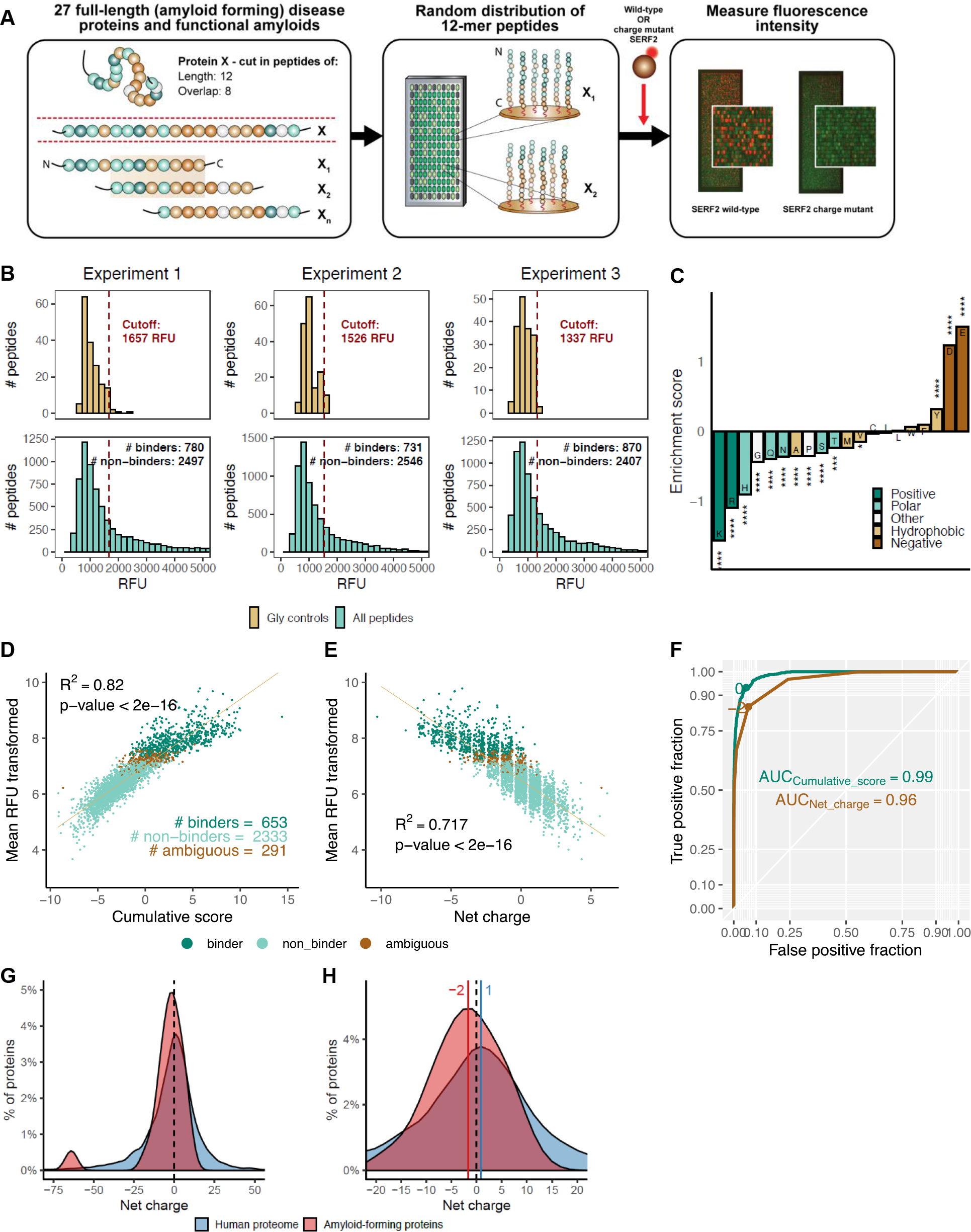
SERF2 interacting peptide sequences are enriched for negatively charged amino acids. (A) Schematic representation of the peptide micro-array screen. Green colour represents autofluorescence of peptides at 532nm. (B) Histograms of the fluorescence intensities of wild-type ATTO633-labeled SERF2, bound to the peptides on the microarray in each of three independent repeats. Fluorescence intensities for Glycine (Gly) controls are marked in yellow (top panels). Red dashed line indicates cutoff between binders and non-binders. The numbers of binders and non-binders identified in each experiment are indicated. Peptides are only classified as binders when both instances of the duplicate has a higher RFU than the cutoff. (C) Enrichment scores (ln(probability ratio)) of all amino acids in SERF2 binding versus SERF2 non-binding peptides. Statistical significance was determined through hypergeometric testing with Bonferroni correction for multiple comparisons. *p<0.05, **p<0.01, ***p<0.001, ****p<0.0001. (D+E) Correlation of the natural logarithm of mean wild-type SERF2 binding intensities (background-corrected per experiment) with the cumulative amino acid enrichment scores (D) or net charge (E) of the micro-array peptides. Mean RFU was transformed as ln(meanRFU – min(mean RFU) + 1). Linear regression curve, R2 and number of binders, non-binders and ambiguous peptides are indicated. (F) Receiver Operating Characteristic (ROC) curves of binder prediction based on cumulative score (blue) or net charge (brown). ROC curve shows fractions of true positive and false positive predictions on y- and x-axes respectively, with increasing cutoff values. Area Under the Curve (AUC) is indicated for both cumulative score and net charge and constitutes a metric for the predictive power of the statistic (cumulative score or net charge). Optimal cutoffs are indicated. (G) Net charge density of amyloid-forming proteins vs. all proteins in the human proteome, represented as the percentage of amyloid-forming proteins or human proteome with a certain net charge. (H) Zoomed-out version of figure 1G, showing the shift of the peak net charges in further detail.

Next, we determined the relative abundance of each individual amino acid in the group of SERF2 binding peptides compared to non-binding peptides. For this comparison, the ratio between the abundance of each amino acid in the group of binding peptides, and the abundance in the group of non-binding peptides was taken, yielding a probability ratio (Dataset S3). We then attributed an enrichment score to each amino acid, by calculating the natural logarithm of the probability ratio (Dataset S3). This score was positive for amino acids that were more abundant, and negative for amino acids that were depleted in the group of SERF2 binding peptides (Figure 1C and Dataset S3). This analysis revealed a more than three-fold overrepresentation of the negatively charged amino acids aspartic acid (Asp, D) and glutamic acid (Glu, E) in SERF2-bound peptides (ln-ratios of 1.23 for Asp and 1.50 for Glu, Figure 1C and Dataset S3). Conversely, when compared to their presence in non-bound peptides, the positively charged amino acids lysine (Lys, K) and arginine (Arg, R) were underrepresented in SERF2-bound peptides (ln-ratios of −1.09 for Arg and −1.56 for Lys, Figure 1C and Dataset S3).

Next we assessed to what extent the cumulative amino acid enrichment scores of the peptides, calculated as the sum of enrichment scores of all amino acids composing that peptide, correlated with their measured SERF2 binding intensities (Figure 1D and Dataset S2). As shown in Figure 1D, the cumulative enrichment scores correlate linearly to the natural logarithm of the actual binding signals (p-value < 2e-16 and R^2^ of 0.82). This observation indicates that a simple scoring function based solely on amino acid composition and completely disregarding position-specific effects, is sufficient to predict binding of SERF2 to peptides. This therefore suggests that SERF2 interaction does not require a strict binding motif. Furthermore, given that charged residues had the most extreme scores in our scoring matrix (negatively charged amino acids scored highest and positively charged amino acids scored lowest), SERF2 binding appears to be mainly driven by net charge. To test this, we performed a linear regression of the natural logarithm of the binding signals versus net charge (Figure 1E). This also showed a strong correlation (p-value < 2e-16), which confirmed that net charge is indeed a key driver for SERF2 binding. However, this regression has a lower R^2^ (0.717) because of a stronger degree of scatter around the regression line, indicating that the cumulative enrichment is a more accurate predictor of SERF2 binding than net charge alone. To confirm this, we plotted Receiver Operating Characteristic (ROC) curves for both predictors (cumulative score and net charge, Figure 1F). In a ROC curve, the fraction of correct binary classifications or “True positive fraction” (SERF2 binder or non-binder) at each threshold of the predictor (either net charge alone or cumulative enrichment score) is plotted against the fraction of false classifications (“False positive fraction”). The Area under the Curve (AUC) gives an indication of the performance of the predictor at the classification problem, in this case classifying peptides into SERF2 binders or non-binders. Although both net charge and cumulative enrichment score show strong predictive power, the cumulative score outperforms net charge (AUC of 0.99 versus 0.96). These results corroborate that net charge seems a key driver of SERF2 interaction, but that likely also non-charged amino acids affect binding intensity.

Since charge seems such an important determinant in SERF2 binding, we asked whether the presence of negative charge is a distinguishing feature of amyloid-forming proteins. We therefore compared the net charges of the amyloidogenic proteins listed in column A of Table S1 with the net charges of all proteins in the full proteome (Dataset S4). By doing so, we found a shift towards more negative net charges (distributed around −2) in the amyloidogenic proteins compared to the rest of the proteome (distributed around +1) (Figure 1G and 1H).

To next profile the SERF2 binding sites to each protein on the microarray, we mapped the binding intensities of each peptide to its corresponding position in its full-length parent protein (Figure 2). Due to the sliding window design of our microarrays, we were able to obtain a SERF2 binding profile by averaging the binding intensities of each of the three peptides that contained a particular residue (Dataset S5). This yielded a binding profile with a resolution of four amino acids (Figure 2 and Figures S1A). To further explore the link between net charge and SERF2 binding, we similarly produced net charge profiles for all the proteins under study (Dataset S5 and Figure S1B), this time averaging the net charges of the three peptides in which a residue is represented (Figure 2 and Figures S1A). This analysis again showed a strong association between SERF2 binding and local negative net charge. Strikingly, while both positively and negatively charged regions are present in the majority of the proteins, all of the proteins analyzed here contain at least one strong negatively charged SERF2 binding site, with the exception of one amyloid-forming protein – human islet amyloid polypeptide (hIAPP) –, the four dipeptide repeat polymers (poly-GR, poly-PR, poly-GA, and poly-PA) and SERF2 itself. Interestingly, despite the strong correlation observed between SERF2 binding and local net charge, we also observed strong variations in binding intensities among peptides with identical net charges (Figures 1E and Figure 2). For example, a clear interaction between SERF2 and a negatively charged region in prion protein around position 150 was observed, while the SERF2 binding intensities were much lower for the two similarly charged, neighboring regions in this protein (Figure 2 and Figure S1A). This again indicated that besides charge, additional sequence properties contributed to the strength of the interactions.

**Figure 2.**
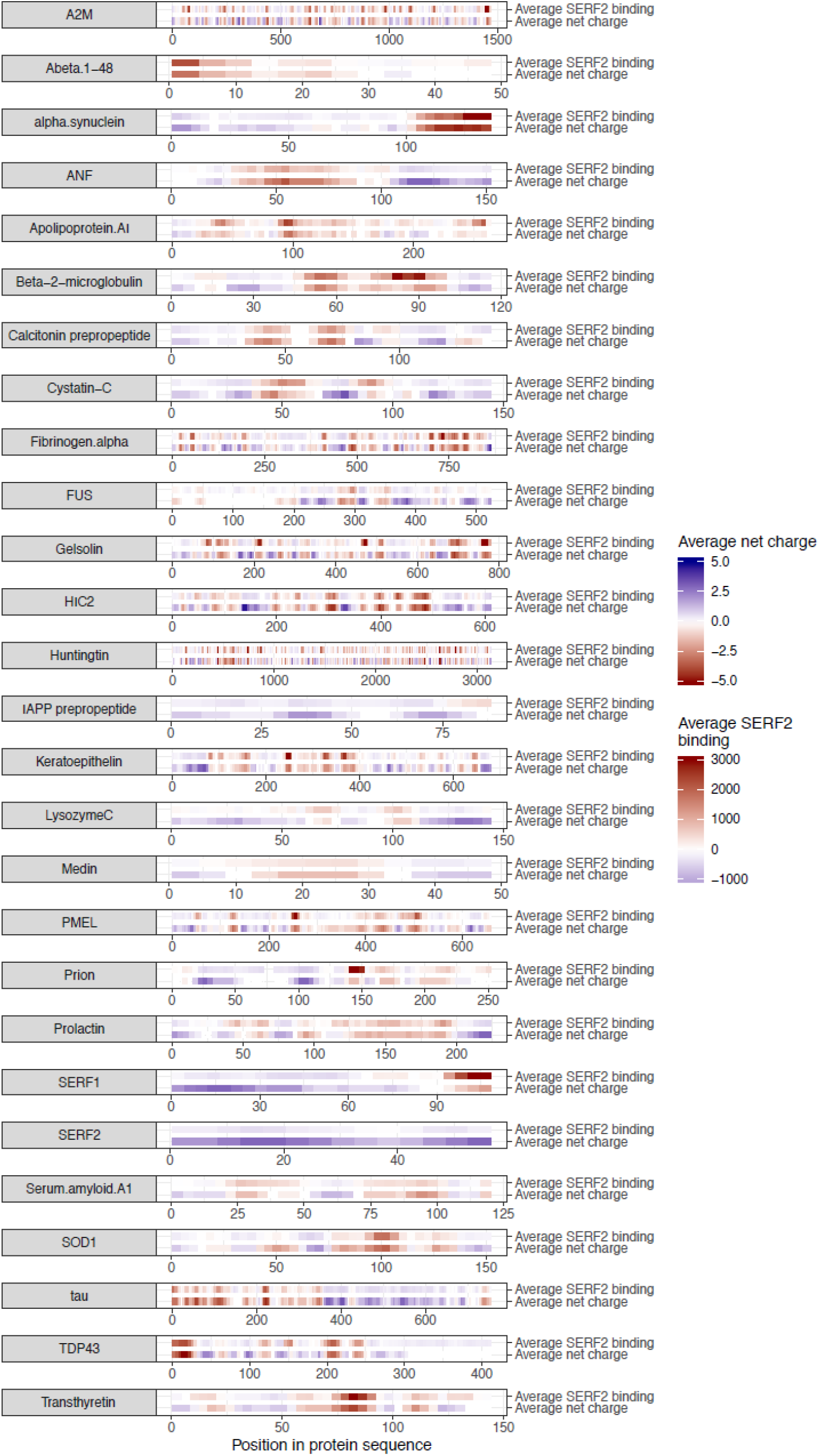
SERF2-binding map to net negatively charged regions. Overview of binding sites for SERF2 in proteins represented on the micro-array slide. Average net charge and binding of SERF2 (see Dataset S5 for values) in each protein is indicated.

### SERF2 binding is further enhanced by hydrophobic, aromatic residues

To identify other sequence features that influenced the interactions of the peptides with SERF2, we focused on the subset of binding peptides for which the actual binding intensities differed strongly (more than two standard deviations over the mean difference) from the predicted intensity based on the cumulative enrichment scores (77 peptides, Figure 3A and Dataset S2). We compared the amino acid composition of these peptides to all other binding peptides (Dataset S6). This analysis revealed a strong enrichment for the hydrophobic, aromatic residues tyrosine (Tyr, Y) and phenylalanine (Phe, F) and the hydrophobic amino acids valine (Val, V) and leucine (Leu, L), suggesting a role for these amino acids in interactions with SERF2 (Figure 3B). Taken together, these findings indicate that the interaction between charged residues and SERF2 is further enhanced by the presence of hydrophobic residues.

**Figure 3.**
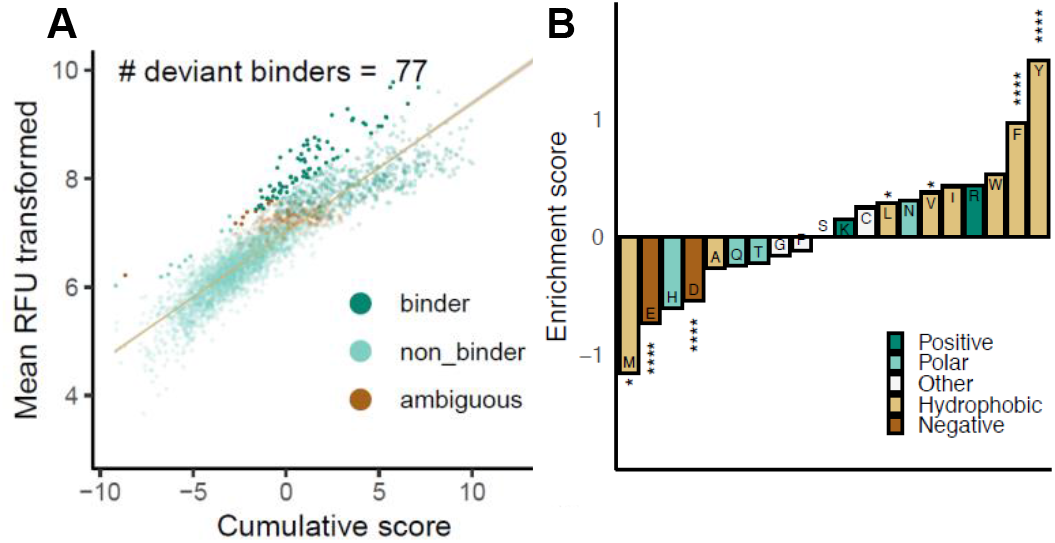
Peptide sequences enriched for hydrophobic, aromatic amino acids show enhanced SERF binding. (A) Reiteration of Figure 1D, with peptides for which the transformed fluorescence intensity deviates strongly from the predicted value indicated. (B) Enrichment scores (ln(probability ratio)) of all amino acids in SERF2-binding peptides for which the transformed fluorescence intensity deviates strongly from the predicted value, versus the remaining SERF2-binding peptides. Statistical significance was determined through hypergeometric testing with Bonferroni correction for multiple comparisons. *p<0.05, **p<0.01, ***p<0.001, ****p<0.0001.

### The positively charged N-terminus of SERF2 mediates binding

SERF2 is a highly positively charged protein that has a net charge of +10, mainly due to a region in its N-terminus that is evolutionarily highly conserved (Figure S2A and S2B). To assess the role of this region in the charge-based interactions between SERF2 and amyloidogenic proteins, we neutralized the net charge of +5 in this region by inducing point mutations in the three positively charged amino acids, Lys^16^, Lys^17^ and Lys^23^ (Figure 4A). To exclude the possibility of the charge mutations resulting in structural changes relative to the wild-type protein, the secondary structure of both proteins was assessed with Fourier-transform infrared spectroscopy (FTIR, Figure S2C). These measurements revealed the structures of both the wild-type and charge mutant SERF2 proteins to be largely disordered: their random coil content was estimated to be over 80% as calculated by peak intensity ratios. This indicates that the charge mutations did not induce major changes in the secondary protein structure (Figure S2C).

**Figure 4.**
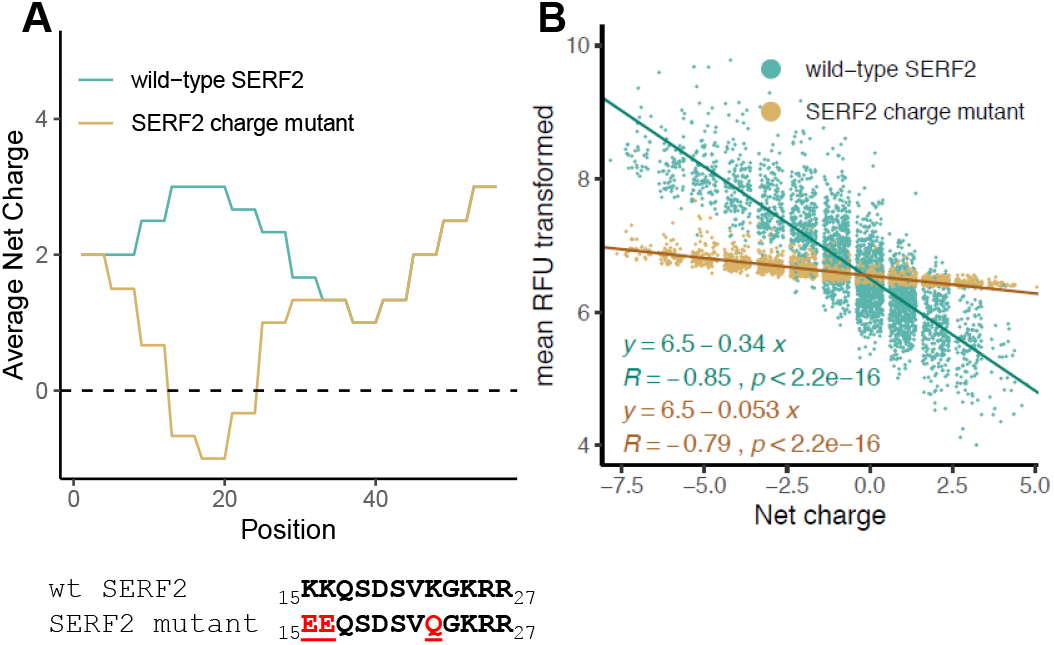
The positively charged N-terminus of SERF2 mediates binding. (A) Mutations and charge distribution in the wild-type SERF2 and SERF2 charge mutant. (B) Correlation between wild-type SERF2 and SERF2 charge mutant binding intensities versus peptide net charge. Mean RFU was transformed as ln(meanRFU – min(mean RFU) + 1). Pearson correlation coefficients (R), p-values and linear regression equations are indicated.

We then tested the ability of the SERF2 charge mutant protein to interact with the peptides on the microarray, and observed that the number of peptides to which it bound (Figure 1A and S2D) was lower than the number of peptides bound by wild-type SERF2 (Figure 1B) in each experiment. Furthermore, the number of peptides to which it bound reproducibly across three experiments (317 peptides) was less than half of the number of wild-type SERF2 bound peptides (653 peptides) (Dataset S2). Although both wild-type and charge mutant SERF2 binding signals showed a significant correlation with net charge, this correlation was much stronger for wild-type SERF2 than for the SERF2 charge mutant (regression slopes of −0.34 and −0.053, respectively, Figure 4B and S2E). The possibility that higher binding intensities were measured for fluorescently labeled wild-type SERF2 than for charge mutant SERF2 due to the presence of multiple fluorescent labels on wild-type SERF2 was excluded by mass spectrometry (Figures S2F-I).

Altogether, these results indicate that charge interactions are a major mediator of the interaction between SERF2 and amyloidogenic proteins, and that the positively charged N-terminus of SERF2 is involved in these interactions. Furthermore, the finding that hydrophobic, aromatic residues were enriched in the strongest SERF2-binding peptides (Figure 3B) suggests that on top of charge-charge interactions, also cation-pi interactions between positively charged residues in the SERF2 N-terminus and the aromatic residues of substrate peptides might mediate SERF2-substrate interactions.

### SERF2 promotes amyloid formation through its positively charged N-terminus

Now our data indicates that charge is an important factor in interactions between amyloidogenic proteins and SERF2, we next aimed to determine the functional consequences of these interactions for amyloid catalysis. Previous work identified the C-terminus of alpha-synuclein as the interaction site for MOAG-4 and SERF1A (11, 13, 14). In line with this observation, our peptide microarray screen revealed that wild-type SERF2 also interacts with the acidic C-terminal region of alpha-synuclein, as well as with the negatively charged N-terminal region of amyloid beta (Figures 5A and 5B). Previous reports have shown that both these regions are important for the solubility of these proteins (17–19). To test whether the binding of SERF2 to the negatively charged regions of alpha-synuclein and amyloid beta promotes amyloid formation of these proteins, we compared the *in vitro* aggregation kinetics of purified alpha-synuclein and amyloid beta in the presence and absence of wild-type or the charge mutant SERF2 using well-established thioflavin T (ThT) fluorescence assays. These experiments showed that the formation of both alpha-synuclein and amyloid beta ThT-positive species was accelerated in the presence of wild-type SERF2 (Figure 5C, 5D and Dataset S7). Equimolar amounts of SERF2 with alpha-synuclein strongly reduced the initial lag phase, and the half-time of conversion was reduced by about 60% (Figure 5C, S3A and Dataset S7). In addition, both the initial lag phase and midpoint of amyloid growth for amyloid beta aggregation were reduced by about 30% in the presence of SERF2 (Figure 5D, S3B and Dataset S7). In contrast, the SERF2 charge mutant did not accelerate the amyloid formation of alpha-synuclein and amyloid beta. These results indicate that the positively charged region of SERF2 is required for its amyloid-promoting effect, which is mediated by the negatively charged regions in amyloidogenic proteins.

**Figure 5.**
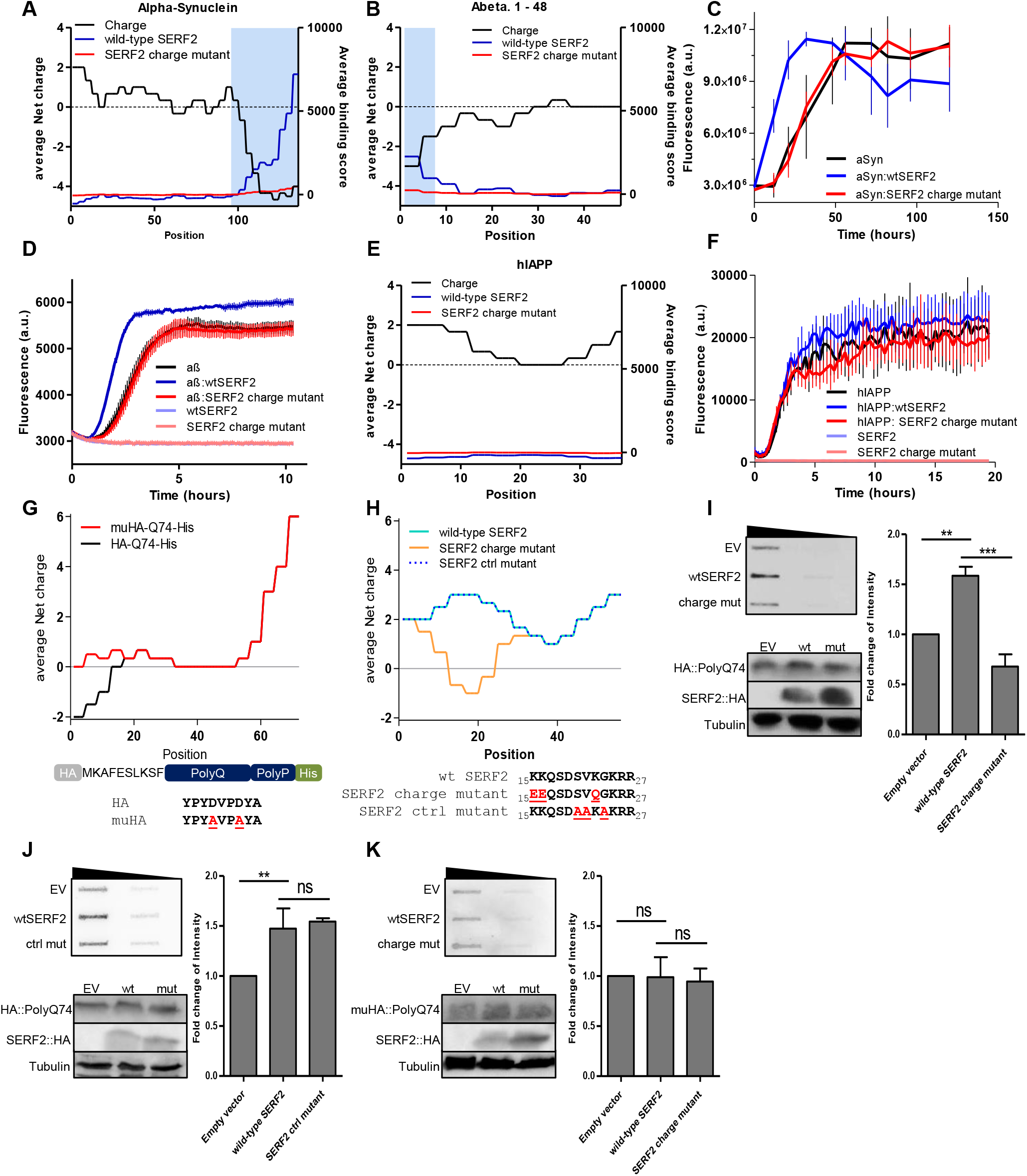
SERF2 drives amyloid formation via charge interactions. (A+B) Net charge distribution (black) of αSyn (A) and Aß (B), plotted together with the binding distribution of wild-type SERF2 (blue) and the SERF2 charge mutant (red). Blue boxes represent the SERF2 interacting regions. (C) ThT-monitored amyloid kinetics of 50uM alpha-synuclein in the presence of either wild-type SERF2 or mutant SERF2 in a 1:1 ratio. The average of three replicates is represented and error bars indicate mean ± SD. Normalized aggregation data in Figure S3A. (D) ThT-monitored amyloid kinetics of 1uM amyloid-beta in the presence of either wild-type SERF2 or mutant SERF2 in a 1:4 ratio. The average of four replicates is represented and error bars indicate mean ± SD. Normalized aggregation data in Figure S3B. (E) Net charge distribution of hIAPP (black), plotted together with the binding distribution of wild-type SERF2 (blue) and the SERF2 charge mutant (red). (F) ThT-monitored amyloid kinetics of 2uM human islet amyloid polypeptide in the presence of either wild-type SERF2 or mutant SERF2 in a 1:4 ratio. The average of five replicates is represented and error bars indicate mean ± SD. Normalized aggregation data in Figure S3C. (G) Model and charge distribution graph of HA-Q74 and muHA-Q74 constructs. (H) Mutations and charge distribution in the SERF2 control mutant and SERF2 charge mutant proteins. (I-K) Filter Trap with 5-fold serial dilution and quantification of crude protein extract from SERF2 CRISPR deletion mutant cells transiently transfected with HA-polyQ74 and either wild-type SERF2, empty vector or SERF2 charge mutant (I), wild-type SERF2, empty vector or SERF2 control mutant (J) or transfected with muHA-Q74 and either wild-type SERF2, empty vector or SERF2 charge mutant. Western blots for HA-Q74, muHA-Q74, SERF2 and Tubulin expression were included as controls (see Figure S3K-M for quantification). For all Filter Trap assays, the depicted blots are from one representative experiment of four biological replicates. The average of four biological replicates with each three or four technical replicates is represented in the graphs. Data are represented as mean ± SD, significance was calculated using a one-way ANOVA, followed by a post-hoc Bonferroni multiple comparisons test. (I) p<0.0001, (J)=0.0003, (K) p=0.9130 **p < 0.01; ***p < 0.001.

To further challenge that SERF2 only has an amyloid-promoting effect if its positively charged region can interact with the amyloidogenic protein, we turned to our peptide microarray screen and selected a peptide for which no interaction with either wild-type SERF2 or the SERF2 charge mutant had been identified, namely hIAPP (Figure 2 and 5E). hIAPP is a predominantly positively charged peptide with strong amyloid-forming properties, which is why we chose this peptide to perform a ThT assay in the presence and absence of wild-type or charge mutant SERF2. This experiment showed that neither wild-type SERF2 nor charge mutant SERF2 had an effect on hIAPP amyloid aggregation (Figure 5F, S3C and Dataset S7). Taken together, these data support our finding that electrostatic interactions between SERF and amyloidogenic proteins are required to allow SERF to catalyze amyloid formation.

### Charge interactions are required for SERF to drive protein aggregation in cells

Our data suggest a role for SERF2 as an amyloid-promoting factor through interactions with negatively charged regions on amyloidogenic proteins. Because negatively charged regions are present in a large fraction of the proteome, we wanted to know whether this mechanism also applies in the context of the full proteome of a human cell. Therefore, we generated a *SERF2* CRISPR-deletion mutant HEK293T cell line (Figure S3D-H) and tested whether adding wild-type and charge mutant SERF2 to these cells affected the aggregation of an aggregation-prone model substrate (Figure 5G). The substrate we chose was an HA-tagged mutant Huntington exon 1 (HTTex1) fragment protein with 74Q repeats. In previous experiments HTTex1 HA-Q74 has shown robust and strong aggregation in cells, which can be modified by several cellular factors, including SERF1A and SERF2 (12, 20). The HA-tag, also found to bind SERF2, adds a net negative charge to the N-terminal site of the mutant HTTex1 fragment protein (Figures 5G and S3I). The HTTex1 HA-Q74 fragment protein was therefore a model protein that combined the elements that we had identified as being essential for interactions with SERF2 with an effective aggregation-prone region. Moreover, this model protein allowed us to determine the contribution of electrostatic interactions between SERF and its client proteins to catalyze protein aggregation in the cell, independent of other cellular factors. Note that we used HTTex1 in this context as a synthetic amyloidogenic polyglutamine peptide, rather than as a model for Huntington’s disease.

We first determined whether the positive charge of SERF2 was required to interact with the HA tag. Therefore, a Filter Trap assay was performed using cell lysates from the SERF2 deletion mutant cell lines that expressed wild-type SERF2, the SERF2 charge mutant or a SERF2 control mutant, in combination with the HTTex1 HA-Q74 fragment protein. A SERF2 control mutant, with substitutions in three uncharged amino acids to alanine (Ala) in the same domain was added to exclude the possibility that mutations in the N-terminal region of SERF2 other than mutations in charged amino acids might be able to diminish the effect of SERF2 on aggregation (Figure 5H). We found that Q74 aggregation was lower when co-expressed with the SERF2 charge mutant (Figure 5I and S3K) than with the wild-type SERF2 or the SERF2 control mutant protein (Figure 5J and S3L). The mutations did not change the localization of the SERF2 charge mutant in the cell when compared with the wild-type protein (Figure S3J). These results confirmed a role for the positively charged N-terminal amino acids of SERF2 in driving aggregation in human cells.

To further establish the role of charge interactions between SERF2 and the HA-Q74 protein, we also generated a mutated version of the HA-Q74 substrate. Mutations in the HA-tag were induced by substituting two negatively charged Asp residues for the neutrally charged amino acid Ala, resulting in a net neutral charge for the HA-tag (Figure 5G). The HA-Q74 and muHA-Q74 constructs were overexpressed in *SERF2* CRISPR-deletion mutant cells in combination with either wild-type SERF2 or the SERF2 charge mutant, and a Filter Trap assay was performed. This experiment revealed that neither wild-type SERF2 nor the SERF2 charge mutant could affect the amount of muHA-Q74 aggregation (Figure 5K and Figure S3M).

The results obtained using this model suggest that SERF2 can trigger aggregation through specific and direct electrostatic interactions with charged regions of aggregation-prone proteins, independently of other modifying factors present in the cell.

### Charge mutations in MOAG-4 reduce aggregation in *C. elegans*

Our *in vitro* results showed that charge mutations in the positively charged N-terminal region of SERF2 could abolish the amyloid-promoting effect of SERF2. We then wanted to determine whether this also applied in the context of a full-body organism. We therefore introduced charge mutations into the *C. elegans* ortholog of SERF2, MOAG-4, to see if this would suppress aggregation in a *C. elegans* model of protein aggregation. We used CRISPR to induce point mutations in the endogenous locus of the *moag-4* gene to change the exact same amino acids as previously done for SERF2 (Figure 6A and S4A). The MOAG-4 charge mutant and MOAG-4 control mutant strains were crossed with a polyglutamine (polyQ) worm model (Q40). Cells in the body-wall muscle of the Q40 worm express a transgene carrying an aggregation-prone polyQ stretch of 40 residues, fused C-terminally to yellow fluorescent protein (YFP, Figure S4B) (12, 21). The effect of the point mutations in MOAG-4 was determined by quantifying the number of aggregates in the worms in the fourth larval stage (L4). As previously reported, genomic deletion of MOAG-4 (MOAG-4 del) strongly reduces the number of aggregates (12). Here we also saw that the number of aggregates in worms carrying mutations in positively charged amino acids at the N-terminus of MOAG-4 was much lower – by about 60% – than the numbers seen in wild-type Q40 worms or worms expressing the MOAG-4 control mutant (Figure 6B, 6C and S4C-ED). Based on the peptide arrays, we do not expect MOAG-4 to bind to the polyglutamine part of Q40-YFP. Whether it interacts with charged residues in flanking regions, as observed for SERF2, or acts indirectly via other molecules in the cell remains to be determined. However, our data reveal that the charge of endogenous MOAG-4 is responsible for its aggregation-promotion in C. elegans as well.

**Figure 6.**
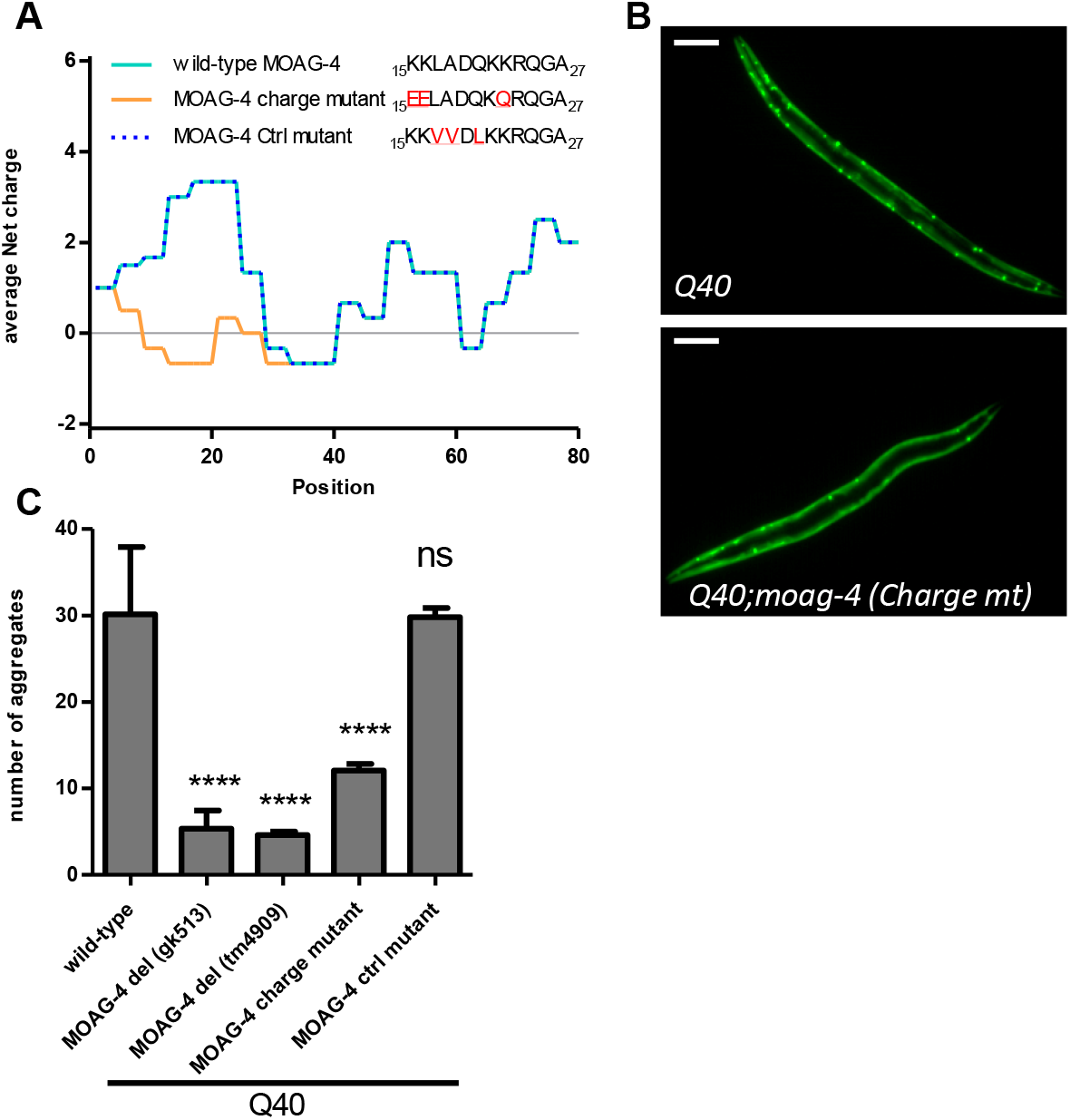
Neutralizing charge of the endogenous SERF ortholog MOAG-4 is sufficient to suppress protein aggregation in C. elegans. (A) Mutations and charge distribution of wild-type MOAG-4, MOAG-4 charge mutant and MOAG-4 control mutant. (B) Representative images of Q40 and Q40;moag-4 charge mutant animals. Scale bar, 75 mm. (C) Representative quantification of the number of aggregates in Q40 worms with moag-4 deletion or expression of either wild-type moag-4, moag-4 charge mutant or moag-4 ctrl mutant. The results shown are representative experiments of three biological replicates of n=20 worms in L4 stage. Data are represented as mean ± SD and significance was calculated using a one-way ANOVA, followed by a post-hoc Bonferroni multiple comparisons test. p<0.0001. ****p < 0.0001.

### Charge rather than amino-acid composition of MOAG-4/SERF drives binding to amyloidogenic proteins and aggregation

Our data indicates that the positive charge in the N-terminus of SERF2 is required for interactions with amyloidogenic proteins and for subsequent acceleration of amyloid formation. To confirm that it is indeed the positively charged nature of Lys^16^, Lys^17^ and Lys^23^ in SERF2 that is responsible for this ability and not any other lysine-specific characteristic, we created a lysine-to-arginine SERF2 mutant (KR-mutant, Figure 7A). The K-to-R mutations do not alter the charge of the SERF2 protein. To investigate if the KR-mutations change the affinity of SERF2 to previously identified strong binders of SERF2 in alpha-synuclein and amyloid-beta (Dataset S2, peptide numbers 61 and 0), we measured their binding affinities using Microscale thermophoresis (MST). MST can determine the binding affinity of a fluorescently labeled molecule to a potential ligand by monitoring the movement of fluorescent molecules through a microscopic temperature gradient. This revealed that at low ionic strength, wild-type SERF2 and the SERF2 KR-mutant have similar binding affinities for the alpha-synuclein peptide (K_D_ = 3uM and 2.6uM respectively, figure 7B and 7D) and for the amyloid-beta peptide (KD = 9.6uM and 6.6uM respectively, figure 7C and 7D). In contrast, the binding affinities for the SERF2 charge mutant with the alpha-synuclein and amyloid-beta peptides are substantially lower (KD = 56uM and 182uM respectively, Figure 7B, 7C and 7D). These data support our hypothesis that charge complementation drives the interactions between SERF2 and substrate proteins, and indicates that no property of lysine, other than its positive charge is required for these interactions.

**Figure 7.**
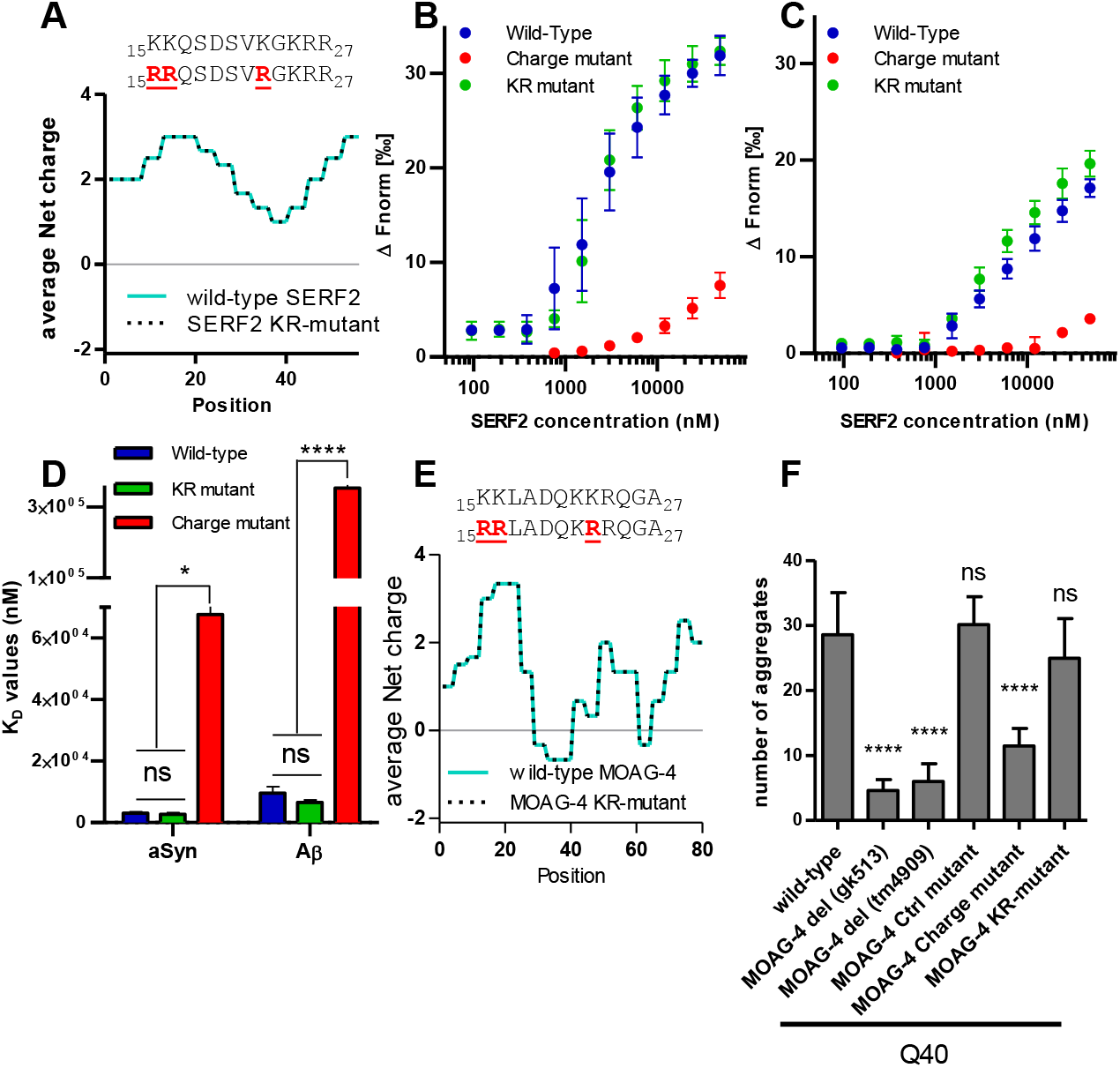
Charge rather than amino-acid composition of MOAG-4/SERF drives binding to amyloidogenic proteins and aggregation. (A) Mutations and charge distribution of wild-type SERF2 and SERF2 KR-mutant. Wild-type SERF2 and KR-mutant sequences are shown, mutations are indicated in red (B+C) Microscale Thermophoresis (MST) analysis of wtSERF2, SERF2 charge mutant and KR-mutant binding to the strongest identified wild-type SERF2-binding peptide from alpha-synuclein (B) and amyloid-beta (C). Binding is represented as the Δ Fnorm, the normalized difference in relative fluorescence between a specific sample and the baseline (the binding intensity without peptide). (D) KD values for each interaction quantified from the experiments depicted in B) and C). n=4. Significance was determined using a two-way ANOVA followed by Tukey’s multiple comparisons test * P-value <0.05, **** P-value < 0.0001. (E) Mutations and charge distribution of wild-type MOAG-4 and MOAG-4 KR-mutant, MOAG-4 sequences are shown and mutations are indicated in red (F) Representative quantification of the number of aggregates in Q40 worms with moag-4 deletion or expression of either wild-type moag-4, moag-4 charge mutant, moag-4 ctrl mutant. Or moag-4 KR-mutant. The results shown are representative experiments of three biological replicates of n=20 worms in L4 stage. Data are represented as mean ± SD and significance was calculated using a one-way ANOVA followed by a post-hoc Bonferroni multiple comarisons test. ****p < 0.0001.

Additionally, we tested if the aggregation-promoting ability of the KR-mutant also remained in our *C. elegans* model for polyQ aggregation. Therefore we crossed a CRISPR induced *moag-4* KR-mutant strain with our Q40 worm model and quantified the number of aggregates in the body-wall muscle of the worms (Figure 7E, 7F and S5A-D). Here we found that the number of aggregates in the worms expressing the KR-mutant was not different from the amount of aggregates found in the body-wall muscle of worms expressing wild-type MOAG-4 (Figure 7F and S5B-D). In summary, these data show that SERF2 and MOAG-4 retain their amyloid-promoting properties when lysines are substituted by arginines and therefore that charge is the driving force for the interactions between SERF2 and amyloidogenic proteins.

## Discussion

While SERF has been identified as an enhancer of protein toxicity and amyloid aggregation for a range of unrelated amyloidogenic proteins, a shared underlying mechanism had not yet been investigated (11–13). Here we found that SERF2 binds preferentially to negatively charged amino acids, and that this binding and its catalyzing effect on amyloid formation required SERF2’s evolutionarily conserved positively charged N-terminal domain. We were particularly excited to find that neutralizing the charge in the endogenous locus of MOAG-4 in *C. elegans* strongly reduced the aggregation of a reporter substrate, which suggests that neutralizing the charge of a single protein in the cell’s proteome is sufficient to alter proteome stability. In the *C. elegans* integrated dataset from the PaxDB (22), MOAG-4 is listed at 387 ppm, which puts it in the top 5% most abundant proteins identified. For SERF2, these levels range from 0.54 till 413, depending on the cell lines in which the concentrations have been measured. In comparison, C. elegans small heat shock protein 12.1 – member of a class of chaperones for which substrate KDs have been reported in the low micromolar range (23–25)– has an abundance of 380 ppm. These comparable levels support the relevance of the low affinity interactions between MOAG/SERF and the substrate proteins that we observed in our cell-free experiments.

Previous studies have described how the processes involved in guiding unfolded proteins towards their properly folded state – and keeping them in this native state – rely on charged residues (26–30). The charge of proteins is affected by numerous factors, including mutations, changes in pH, post-translational modifications and external stress (31–33). In addition, other modifying factors, such as charged polymers, are known to promote amyloid fibril formation. For example, polyanions like glucosaminoglycans (i.e. heparin), nucleic acids, and polyphosphate (polyP), as well as positively charged polylysines and polyamines were recognized to interact with oppositely charged regions, compensating their charge and promoting amyloid formation (17, 34–38). These factors can thereby lead to excessive unfolding and exposure of unprotected aggregation-prone regions. When the abundance of unfolded proteins exceeds the capacity of the cellular protection mechanisms, these unfolded proteins become susceptible to off-pathway structural conversions that drive them into thermodynamically highly stable amyloid fibrils (39–42). Our findings suggest that cellular modifiers such as SERF may similarly accelerate protein transitions to amyloid by acting on the protein’s charge.

Most of what is known about SERF’s mechanism of action comes from studies on alpha-synuclein. Alpha-synuclein is a natively unstructured protein that remains soluble through a strong interaction between its acidic C-terminus and its N-terminal region, an interaction that shields its aggregation-prone middle region (17, 18, 43). Factors that can disrupt these electrostatic intermolecular interactions, such as high salt concentrations (11), polyamine compounds (17), metal cations (44) or C-terminal truncations (13, 45), have been shown to enhance alpha-synuclein aggregation. In previous Nuclear magnetic resonance spectroscopy studies, SERF1A or MOAG-4 was found to interact with the amino acids in the C-terminal region of alpha-synuclein (11, 13). This interaction exposes an amyloid nucleation site on alpha-synuclein that is otherwise concealed by intermolecular interactions (14). A direct but transient interaction between SERF and alpha-synuclein has been confirmed by overexpression of SERF1A and alpha-synuclein in SHSY-5Y neuroblastoma cells (14). Interestingly, the interactions between alpha-synuclein and SERF1A or its yeast ortholog ScSERF have been shown to result in the formation of a fully disordered protein complex, which accelerates the primary nucleation of alpha-synuclein amyloid formation (14, 46, 47). Our findings are in line with these observations and suggest that also SERF2 might accelerate primary nucleation of alpha-synuclein amyloid formation through charge complementation.

Here we observed a similar effect of SERF2 on amyloid beta. In amyloid beta, the strongest negative charge is located in the N-terminal region of the protein. Interaction of SERF2 with amyloid beta appears to take place at this charged region: when we added SERF2 to amyloid beta in kinetic assays we saw that SERF2 required this charge interaction to accelerate aggregation. Recent studies have shown that it is the N-terminus of the amyloid beta peptide that determines its tendency to aggregate. Indeed, N-terminal mutations that affect the charge of this region are known to modify amyloid beta fibril formation. Such mutations include amino acid changes (48, 49), post-translational modifications (50) and truncations of the protein (19, 51–53). In addition, a number of residues in the N-terminus of amyloid beta can bind metal cations – including Cu^2+^ and Zn^2+^ – which also accelerate amyloid beta aggregation through mechanisms that are currently unresolved (54–57). Given the fact that charge is a common denominator in all of these modifying factors, mechanisms similar to those observed for MOAG-4/SERF may well explain their amyloid-promoting effect.

As mentioned above, our findings are in line with the previous binding site for the aggregation-promoting effects of MOAG-4 and SERF1A on alpha-synuclein (11, 14). However, besides the previously proposed mechanism that focuses on the specific disruption of protective intermolecular interactions in alpha-synuclein, also a more general mechanism that involves the stability of any aggregation-prone protein may apply. Next to hydrophobicity and beta sheet propensity, net charge is a critical determinant for the behavior of proteins in solution, because charge repulsion between molecules is essential to keep proteins uniformly distributed. This phenomenon is known as colloidal stability, and strongly determined by the charge distribution of the molecules (58, 59). Neutralization of charged regions – as occurs in the presence of mutations, modifications or modifiers such as SERF – could result in reduced charge repulsion between amyloid proteins and lower the colloidal stability of the proteins in solution, thus accelerating fibril formation. Colloidal stability has also been shown to be important for liquid–liquid phase separation, where proteins are present in high concentrations in membrane-less cellular compartments (60). Mutations, post-translational modifications or environmental factors that result in imbalances in colloidal stability – including charge alteration – could initiate a liquid-to-solid transition towards aggregate-like structures, a transition that resembles the process of amyloid formation (61–63). A third possibility is the existence of other activities of SERF2 that are initiated by electrostatic interactions between SERF2 and amyloid-forming proteins. Additional research is required to distinguish which mechanisms apply for SERF2 in the presence of amyloidogenic proteins.

Why SERF acts as an amyloid-promoting factor is not known. Misfolded monomeric and oligomeric species are generally considered to be the toxic species in age-related disease. A low-energy sequestration mechanism that quickly removes and compactly stores unwanted aggregation-prone proteins could therefore be beneficial. It is possible that under healthy conditions SERF’s function is part of such a sequestration mechanism, whereby it drives transitions of aggregation-prone proteins toward an amyloid state. Alternatively, the aggregation-promoting effect of MOAG-4/SERF2 in the presence of aggregation-prone disease proteins could be an unwanted side effect of a different function. For example, a role for SERF1A as an RNA chaperone in the formation of liquid-like RNA organelles has been suggested (16). Since both RNA and alpha-synuclein are known to interact with the positively charged N-terminal region of SERF, they might compete for this SERF binding site, possibly favoring alpha-synuclein–SERF interactions and additional amyloid formation under stress conditions (16). Meyer et al’s findings, together with the previously shown toxic effects of SERF2 in the presence of amyloid proteins (12), could suggest that the amyloid-promoting properties of SERF2 are a function that is induced under stress conditions.

Based on our results and those of recently published studies on SERF1A and ScSERF, we propose a mechanistic model in which charge complementation by SERF accelerates the primary nucleation of amyloid. Our finding that simply changing the charge of the endogenous SERF in C. elegans was sufficient to have a profound effect on protein aggregation indicates a critical role for charge complementation in the regulation of proteome stability. Taken together, our results suggest that preventing charge interactions between aggregation-prone proteins and charged cellular modifiers deserves exploration as a strategy to prevent or delay the onset of protein toxicity in ageing and age-related diseases.

## Materials and methods

### Microarray peptide screen

Custom-made microarrays were purchased from PEPperPRINT (PEPperCHIP® Peptide Microarray, PEPperPRINT). For the production of the microarrays, 12-mer peptides were directly synthesized on poly(ethylene glycol)-based graft copolymer-coated glass slides with a three amino acid linker (ß-alanine, aspartic acid, ß-alanine). All experiments were performed using the HS 400™ Pro Hybridization Station and HS Pro control manager software (Tecan). Slides were incubated with 1µM ATTO633-labeled wild-type or charge mutant SERF2 proteins in binding buffer, and imaged and analyzed using the Powerscanner Microarray and Array-Pro® Analyzer software (Tecan) with wavelengths channel 1: 580/30 nm and channel 2: 676/37 nm.

### Microarray data analysis

All peptides on the microarray were classified as SERF2 “binders”, “non-binders” or “ambiguous” as follows. For each experiment, a cutoff value distinguishing binder from non-binder peptides was calculated as the mean relative fluorescence units (RFU) of the Gly control peptides plus two times the standard deviation of the RFU signals of the Gly control peptides, as shown in Figure 1B. Assuming an approximately normal distribution for the binding signal to Gly control peptides, this cutoff would encompass 97.5 % of the control peptide population, indicating that peptides with higher RFU signals likely do not belong to the background population, and are in fact true binders. Per experiment, a peptide was classified as a binder if both of its duplicates on the microarray showed higher binding intensities than the cutoff value (Dataset S1). Peptides for which this was not the case were considered non-binders. Finally, only peptides that fulfilled the binder criterium in each of the three repeat experiments were considered actual binders. Peptides that consistently fell below the cutoff value across three experiments were classified as non-binders. The remaining peptides, which showed an inconsistent classification across three experiments, were classified as “ambiguous” (Dataset S2).

Next, amino acid enrichment scores – also referred to as ln(probability ratios) – in binding versus non-binding peptides were determined using the formula: 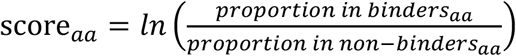, with aa indicating each of the 20 amino acids (Dataset S3). Ambiguous peptides were not considered in these analyses. Statistical significance of these enrichments was determined through hypergeometric testing with Bonferroni correction for multiple comparisons. Using the enrichment scores for each amino acid, we then produced a cumulative score for each peptide using the formula:

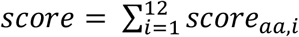 with score_aa,i_ indicating the score for the amino acid aa in position I of the peptide (Dataset S2). The performance of this cumulative score as a predictor for SERF2 binding was assessed through linear regression and ROC curve analysis as indicated in the main text.

To assess how amino acid composition differs between binders that deviate strongly from their predicted value based on the cumulative scores and binders that do not, we identified a subset of binders as “deviating” if their mean RFU values were more than two standard devations above the mean difference with the regression curve across all data points (as shown in Figure 3A). We then repeated the workflow above to identify enrichment scores, this time calculating deviant scores as follows: deviantscore 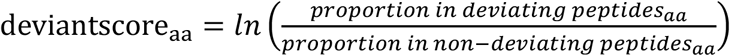 with aa indicating each of the 20 amino acids (Dataset S6).

For an overview of SERF2 binding sites along the primary sequence of the proteins represented on the microarray, the average binding intensity for the Gly control peptides was first subtracted from the binding signals in each repeat experiment, yielding background-corrected binding signals. Next, for each residue, the average net charge and average binding intensity was calculated by taking the average charge or intensity of all three 12-mer windows in which this residue was represented. This resulted in average-score patterns with a resolution of four amino acids as depicted in Figure 2 and Supplemental Figure S1B (Dataset S5).

The analyses above were performed using a combination of Python 3.6 (PyCharm IDE version 2018.1.2) and R statistical software version 3.5.2 (IDE RStudio version 1.1.463).

### Microscale thermophoresis (MST)

Peptides were synthesized in-house using an Intavis Multipep RSi synthesizer following Fmoc/tBu solid-phase synthesis strategy. Peptides were subsequently labeled with 5(6)-carboxyfluorescein as described in the supplementary information.

Labeled peptides were diluted to a concentration of 600 nM in 10 mM Tris pH 7.6 with 0.5% Tween-20. SERF2 or mutants thereof were diluted to a concentration of 107 µM in 10 mM Tris, and a 1:1 dilution series produced. This dilution series was then mixed at a 9:1 ratio with 600nm peptideThermophoresis was then performed on a Monolith NT. Automated (NanoTemper, Germany). KD determination was performed using the MO.Affinity Analysis (NanoTemper, Germany) software version 2.3, using a KD model and keeping the target concentration constant at 60 nM.

More details about procedures and reagents used in this study, including descriptions of standard procedures to generate and analyze cell lines, C.elegans strains, and purified proteins, and descriptions of *in vitro* aggregation assays, amyloid-forming proteins versus proteome net charge analysis, and filter trap analyses are provided in the *Supplementary Information, Materials and Methods*.

## Author contributions

A.P, J.S., F.R. and E.A.A.N conceived and designed the research; A.P. and B.H. designed, performed and analyzed the experiments with help from L.J., W.H., A.M.C., M.d.V., R.G., M.K., E.S., M.d.V. and S.L.E.; A.P. and R.G. designed and performed the microarray experiments with help from B.H., J.S., F.R. and E.A.A.N.; B.H. analyzed the data with help from J.S., F.R., A.P. and E.A.A.N; F.A.A. and F.S.F. performed in vitro ThT kinetic assays; R.S. performed *C. elegans* experiments; M.K. designed the illustrations; M.V. provided invaluable mentorship and guidance to F.A.A.; A.P., B.H. and E.A.A.N wrote the manuscript with input from all authors.

## Acknowledgments

We thank the Caenorhabditis Genetics Centre (funded by the NIH National Centre for Research Resources and the NIH Office of Research Infrastructure Programs [P40 OD010440]), and the Mitani laboratory for the *C. elegans* strains (funded by the Japan National BioResource Project). Part of the work was performed in the UMCG Microscopy and Imaging Center (UMIC). We thank Marcel de Vries for help with mass spectrometry (Interfacultair Massaspectrometriecentrum RUG, UMCG), and Sally Hill for critical comments and for editing the manuscript. This project was funded by a Meervoud Grant from NWO (836.09.001) (to E.A.A.N.), a European Research Council (ERC) starting grant (281622 PDControl) (to E.A.A.N.), the Alumni chapter Gooische Groningers facilitated by the Ubbo Emmius Fonds (to E.A.A.N), an Aspasia fellowship from NWO (015.014.005) (to E.A.A.N.), a Boehringer Ingelheim Fonds travel grant (to A.P.), a Cornelis de Cock grant (to A.P.), a Marie Curie Fellowship (to A.M.C.) and a BCN-BRAIN grant (to M.K.). The Monolith NT automated (NanoTemper) instrument was funded by the Funds for Scientific Research Flanders (FWO; Hercules Foundation grant AKUL/15/34 - G0H1716N). The Switch Laboratory was supported by grants from the Flanders Institute for Biotechnology (VIB; grant no. C0401), the Industrial Research Fund of KU Leuven (“Industrieel Onderzoeksfonds”) and the Funds for Scientific Research Flanders (FWO; Hercules Foundation grant AKUL/15/34 - G0H1716N). B.H. was supported by a PhD Fellowship from the IWT (file nr.141546). F.A.A. was supported by a Senior Research Fellowship award from the Alzheimer’s Society, UK (grant number 317, AS-SF-16-003).

## Contact for Reagent and Resource Sharing

Further information and requests for resources and reagents should be directed to and will be fulfilled by Ellen A.A. Nollen. Published research materials and reagents from ERIBA are shared with the academic community under a Material Transfer Agreement (MTA).

## Notes

https://unishare.nl/index.php/s/ggoGMt6Jdo85RFk

### Competing Interest Statement

The authors have declared no competing interest.

