## Supplemental Information for "The cellular modifier MOAG-4/SERF drives amyloid formation through charge complementation"

**This PDF file includes:**

SI Materials and Methods

SI Materials table

SI References

SI Figures S1-S5

Supplementary Table S1, datasets S1-S7 and the R-scripts corresponding to this manuscript can be accessed through: https://unishare.nl/index.php/s/ggoGMt6Jdo85RFk

#### SI Materials and methods

##### Plasmids and constructs

Wild-type *SERF2* DNA (Uniprot identifier P84101-1, length 59 aa) was inserted in the pGEX-4t-1 vector using primers pGEX primer_F1 and pGEX primer_R1, and restriction enzymes EcoRI and NotI. For the generation of the SERF2 charge mutant pGEX-4t-1 construct, the Q5® Site-Directed Mutagenesis Kit (NEB; E0554S) was used according to the manufacturer's protocol, using SDM primer_F1 and SDM primer_R1, and pGEX-4t-1 wild-type *SERF2* construct as a template. Primers were designed using NEBaseChanger™. For mammalian expression vectors, the Q5® Site-Directed Mutagenesis Kit (NEB; E0554S) was used according to the manufacturer's protocol, using a pcDNA3.1-vector containing wild-type human *SERF2* DNA C-terminally fused to an HA-tag as a template. SDM primer_F1 and SDM primer_R1 were used for the SERF2 charge mutant and SDM primer_F2 and SDM primer_R2 for the SERF2 control mutant. The constructs were transformed into NEB 5-alpha Competent *E. coli* (NEB; C2987H) and plated on selection plates containing ampicillin (100ug/mL). The presence of the mutations was confirmed by sequencing. For the generation of the SERF2 CRISPR-deletion mutant HEK293T cells, the pX458 vector (Addgene #48138) was digested using BbsI (NEB), and 100 µM oligos for each gene were annealed using T4 PNK (NEB) following the manufacturer's protocol. 100 ng restricted plasmid and annealed oligos (1/200) were ligated using T4 DNA ligase according to the manufacturer's protocol. Oligos for SERF2 are Serf2_CRISPR_F and Serf2_CRISPR_R.

##### Protein purification

Recombinant wild-type, charge mutant SERF2 and SERF2 KR-mutant fused to glutathione S-transferase (GST) were expressed and purified using the pGEX-4t-1 vector according to standard purification methods.  in *E. coli* competent BL21 (DE3) cells (Promega) and grown overnight at 37°C in 50mL LB medium supplemented with 100ug/mL ampicillin under constant shaking at 220rpm. Expression of the genes was initiated using 1mM IPTG (Promega, V3951). Cells were pelleted by centrifugation, resuspended in PBS supplemented with protease inhibitor cocktail tablets (25x) (Roche; 11697498001) and frozen at -20°C. For protein purification, the cells were lysed by sonication. To remove RNA/DNA, the sonicated cells were supplemented with 0.5M $MgCl_2$ to a final concentration of 6mM and incubated with 5ul/mL Benzonase (Merck Millipore, 70746) at 4°C for 1 hour. Cell debris was removed by centrifugation at 20,000rpm for 40 min at 4°C. The supernatant was filtered using a 0.45 µM filter (VWR; 514-0063) and loaded onto a GSTrap 4B column (GE Healthcare, 28-4017-45) that had been pre-equilibrated with PBS. The column was washed with PBS and the proteins were cleaved from the GST-tag using 80U of Thrombin Protease (GE Healthcare, 27-0846-01), overnight (12–16hr) at room temperature (22-25°C). The proteins were eluted in PBS. For elution, the GSTrap 4B column was attached on top of a HiTrap Benzamidine FF column (GE Healthcare, 17-5143-02), pre-equilibrated with PBS, to remove Thrombin protease from the cleaved off proteins. The purity of the eluted sample was determined with SDS-PAGE followed by Coomassie staining. The presence of full-length wild-type or charge mutant SERF2 was determined by Western blot and mass spectrometry analysis. As SERF2 does not contain any aromatic amino acid residues, the protein concentrations were quantified using the $A_{205}$ customized method on a NanoDrop spectrophotometer for protein and peptide quantification with extinction coefficient 27.

For the in vitro aggregation experiments, amyloid beta and hIAPP were prepared as previously described (1, 2).

**Protein labeling and protein preparation**

For visualization of wild-type or charge mutant SERF2 during SERF2-peptide binding determination, proteins were labeled with an amine-reactive ATTO633 label (ATTO 633 NHS-ester 1 mg, ATTO-tec GmbH, #AD633-31) which can be detected in the red spectral region. ATTO633 NHS-ester stock was prepared in DMSO to a concentration of 1mM. After 10x dilution of the dye in binding buffer (1x PBS, 0.05% Tween, pH 7.4) to a concentration of 100uM, the dye was added to the unlabeled wild-type or charge mutant SERF2 proteins in a 1:1 ratio and incubated in the dark for 1hr on ice. Unbound ATTO633 dye was separated from the labeled proteins using a Sephadex G-25 column (GE Healthcare; 17-0851-01) that had been equilibrated in binding buffer. Proteins were eluted from the column using binding buffer. Labeled protein concentrations were determined using the ‘Proteins&Labels’ application of the NanoDrop spectrophotometer, and by comparing labeled SERF2 proteins with unlabeled SERF2 proteins of known concentrations on Western blot. Blots were quantified using Image Studio Lite software, Version 5.2. The absence of wild-type or charge mutant SERF2 with multiple ATTO633 labels was confirmed using mass spectrometry analysis.

**Microarray peptide screen**

For SERF2-peptide binding determination, custom-made microarrays were purchased from PEPperPRINT (PEPperCHIP® Peptide Microarray, PEPperPRINT). For the production of the microarrays, 12-mer peptides were directly synthesized on poly(ethylene glycol)-based graft copolymer-coated glass slides with a three amino acid linker (ß-alanine, aspartic acid, ß-alanine), using an established peptide laser printing technology (3). Peptides were synthesized from C- to N-terminus, yielding peptides with free N-termini, and C-termini coupled to the slide through the aforementioned linker. In brief, each activated amino acid, covered at its N-terminus by a 9-fluorenylmethoxycarbonyl (Fmoc) protection group, was separately attached with its C-terminus to the three amino acid linker or its neighboring amino acid through a cycle composed of a melting (coupling) step, removal of excessive monomers and removal of the Fmoc protection group. This cycle was repeated until all peptides were synthesized. All 12-mer peptides were present in duplicate and randomly distributed on each half of a slide.

All experiments were performed using the HS 400™ Pro Hybridization Station and HS Pro control manager software (Tecan). The slides were washed with MilliQ three times 1 minute with 30 seconds soak time, followed by washing with binding buffer (1x PBS, 0.05% Tween, pH 7.4) three times 1 minute with 30 seconds soak time. Slides were blocked with blocking buffer (1% BSA, 1x PBS, 0.05% Tween, pH 7.4) for 30 minutes with medium agitation frequency. Slides were incubated with 1µM ATTO633-labeled wild-type or charge mutant SERF2 proteins in binding buffer for one hour with medium agitation frequency. After incubation, the slides were washed with washing buffer (1x PBS, 0.1% Tween, pH 7.4) three times two minutes with 30 seconds soak time, followed by washing with binding buffer three times two minutes with 30 seconds soak time. The slides were then washed with MilliQ three times 1 minute with 30 seconds soak time, and dried with nitrogen for two minutes. The viscosity during all steps was set to medium and all steps were performed at 25°C. The slides were imaged and analyzed using the Powerscanner Microarray and Array-Pro® Analyzer software (Tecan) with wavelengths channel 1: 580/30 nm and channel 2: 676/37 nm, 200% gain and 1% laser power. Slides were regenerated in between experiments. For regeneration, slides were incubated in regeneration buffer (100mM glycine, 500mM NaCl, 6M Gua-HCl, pH 2.0) while sonicated in a water bath at room temperature for 90 minutes. The slides were left overnight in regeneration buffer in the water bath at room temperature. Subsequently, slides were sonicated again for 30 minutes at 50°C. After sonication, the slides were washed five times with MilliQ, five times with absolute ethanol and three times with acetone. Slides were dried using air flow and imaged using the Powerscanner Microarray as described above to confirm the slides were fully regenerated.

**Microarray data analysis**

All peptides on the microarray were classified as SERF2 "binders", "non-binders" or "ambiguous" as follows. For each experiment, a cutoff value distinguishing binder from non-binder peptides was calculated as the mean RFU of the Gly control peptides plus two times the standard deviation of the RFU signals of the Gly control peptides, as shown in Figure 1B. Assuming an approximately normal distribution for the binding signal to Gly control peptides, this cutoff would encompass 97.5 % of the control peptide population, indicating that peptides with higher RFU signals likely do not belong to the background population, and are in fact true binders. Per experiment, a peptide was classified as a binder for that experiment if both of its duplicates on the microarray showed higher binding intensities than the cutoff value (Dataset S1). Peptides for which this was not the case were considered non-binders for that experiment. Finally, only peptides that fulfilled the binder criterium in each of the three repeat experiments were withheld as actual binders. Peptides that consistently fell below the cutoff value across three experiments were classified as non-binders. The remaining peptides, which show an inconsistent classification across three experiments, were classified as “ambiguous” (Dataset S2).

Next, amino acid enrichment scores – also referred to as ln(probability ratios) – in binding versus non-binding peptides were determined using the formula:

$\text{score}_{aa} = ln\left(\frac{proportion\ in\ binders_{aa}}{proportion\ in\ non{-}binders_{aa}}\right)$, with aa indicating each of the 20 amino acids (Dataset S3). Ambiguous peptides were not considered in these analyses. Statistical significance of these enrichments was determined through hypergeometric testing with Bonferroni correction for multiple comparisons. Using the enrichment scores for each amino acid, we then produced a cumulative score for each peptide using the formula:

$score = \sum_{i=1}^{12} score_{aa,i}$ with $\text{score}_{\text{aa,i}}$ indicating the score for the amino acid aa in position i of the peptide (Dataset S2). The performance of this cumulative score as a predictor for SERF2 binding was assessed through linear regression and ROC curve analysis as indicated in the main text.

To assess how amino acid composition differs between binders that deviate strongly from their predicted value based on the cumulative scores and binders that do not, we identified a subset of binders as “deviating” if their mean RFU values were more than two standard devations above the mean difference with the regression curve across all data points (as shown in Figure 3A). We then repeated the workflow above to identify enrichment scores, this time calculating deviant scores as follows: $\text{deviantscore}_{\text{aa}} = ln\left(\frac{proportion\ in\ deviating\ peptides_{aa}}{proportion\ in\ non-deviating\ peptides_{aa}}\right)$ with aa indicating each of the 20 amino acids (Dataset S6).

For an overview of SERF2 binding sites along the primary sequence of the proteins represented on the microarray, the average binding intensity for the Gly control peptides was first subtracted from the binding signals in each repeat experiment, yielding background-corrected binding signals. Next, for each residue, the average net charge and average binding intensity was calculated by taking the average charge or intensity of all three 12-mer windows in which this residue was represented. This resulted in average-score patterns with a resolution of four amino acids as depicted in Figure 2 and Supplemental Figure S1B (Dataset S5).

The analyses above were performed using a combination of Python 3.6 (PyCharm IDE version 2018.1.2) and R statistical software version 3.5.2 (IDE RStudio version 1.1.463).

**Amyloid-forming proteins versus proteome net charge analysis**

Protein sequences for the entire human proteome were obtained from the Uniprot Knowledgebase (UniProt Consortium, 2018; reference proteome UP000005640). The group of amyloid-forming proteins was defined as described in the main text (amyloidogenic proteins listed in Table S1). For all sequences, net charges were calculated as the sum of the numbers of Arg and Lys minus the sum of the numbers of Asp and Glu in the primary protein sequences (Dataset S4). Density plots were generated using R statistical software version 3.5.2 and its IDE RSstudio version 1.1.463.

**Fourier-transform infrared spectroscopy (FTIR)**

FTIR spectra for SERF2 and the SERF2 charge mutant were measured in a Tensor 27 FTIR spectrometer (Bruker) according to the manufacturer's instructions. Both proteins were measured at a final concentration of 1.3 mg/mL in PBS. PBS without protein was measured as blank to produce a background spectrum. Amide I Peaks were deconvoluted using OPUS software in order to determine secondary structure content. Briefly, background spectra were subtracted from the sample spectra, after which the second derivative of the amide I peak was calculated in order to determine the approximate number of peaks to take into the deconvolution step (three peaks for both SERF2 and SERF2 mutant). Next, the amide I peak was deconvoluted with starting peak maxima at 1627 (β-sheet), 1650 (random coil) and 1680 $cm^{-1}$ (β-turn). Peak shapes were set to Lorentz, and widths and intensities were optimized using the local least squares method. As this resulted in peak intensities of 0 for the 1680 $cm^{-1}$ peak in both SERF2 and the SERF2 charge mutant, these were omitted and the remaining peaks were refitted using the local least squares method. Secondary structure content was then calculated as the ratio between the intensity of each peak and the total intensity of all peaks.

***In vitro* aggregation assay**

Alpha-synuclein thioflavin T (ThT) fluorescence measurements were performed in triplicate as described previously (5). In brief, 50 µM alpha-synuclein was mixed with an equimolar amount of SERF2 or SERF2 charge mutant in 50mM Tris-HCl, 150mM NaCl and 3mM $NaN_3$ (alpha-synuclein working buffer, pH 7.4), and shaken at 1,400 rpm at 37°C. At the different time points, 5µl aliquots were removed and diluted in 1mL of a 5µM ThT solution buffer (Sigma-Aldrich), and ThT fluorescence was measured on a Jasco FP-6500 fluorescence spectrometer (Jasco GmbH, Pfungstadt, Germany) at 25°C and 482nm, upon excitation at 442nm, in a 1mL quartz cuvette at 25°C. Ex/em slit widths = 10/10nm.

Aggregation kinetic assays for amyloid beta and hIAPP were performed as described previously (1, 2). In brief, all measurements were performed in black, low-binding 96-well half-area plates, with clear bottom (Corning). Plates were sealed and ThT fluorescence was measured at each time point with an excitation wavelength of 440nm and an emission wavelength of 480nm at 37°C, using a CLARIOstar plate reader (BMG Labtech). Measurements were performed in quadruplicate and quintuplicate, without agitation. 1µM amyloid beta was mixed with 4µM SERF2 or SERF2 charge mutant in 20mM sodium phosphate buffer pH 8.0, 200uM EDTA and 0.02% $NaN_3$, and supplemented with 20uM ThT. 2µM hIAPP was mixed with 8µM SERF2 or SERF2 charge mutant protein in standard PBS.

**Cell lines and culture**

Wild-type HEK293T and SERF2 CRISPR-deletion mutant cells were cultured in Dulbecco's Modified Eagle Medium (DMEM, Gibco; high glucose, pyruvate, 41966052), supplemented with 10% bovine cow serum (BCS; Sigma 12133C) and 1% penicillin/streptomycin (Gibco; 10,000 U/mL, 15140122), at 37°C, 5% $CO_2$. For passaging of cells, 0.05% Trypsin-EDTA (1X), Phenol Red (Invitrogen; 25300-054) was used. Regular mycoplasma tests were performed.

**Generation of SERF2 CRISPR-deletion mutant cell lines**

The SERF2 CRISPR-deletion mutant cell lines were generated as described by Ran *et al.* 2013 (Figure S3D-E)(6). Briefly, cells were transfected using 1mg/mL polyethylenimine (PEI) reagent (Polysciences, 239662) and 500 ng DNA of the pX458-edited plasmid per 200,000 cells, and plated in a 6-well plate. Twenty-four hours after transfection, the cells were trypsinized (Gibco; 0.05% Trypsin-EDTA (1x), Phenol Red, 25300-054) and collected. The cells were diluted to an end concentration of 10 cells/mL in 40% conditioned medium, and 100μL was transferred to each well of a 96-well plate. After 24 hours, wells with GFP-positive cells were selected and grown for 1–2 weeks before being transferred to a 24-well plate. Medium was refreshed every 3–4 days. Mutations in both SERF2 alleles were confirmed with sequencing (Figure S3F), and absence of SERF2 was confirmed by qPCR and Western blot analysis (Figure S3G and S3H).

**Quantitative PCR**

Total RNA was extracted from cell pellets using TRIzol Reagent (Life Technologies) according to the manufacturer's protocol. Total RNA quality and concentrations were assessed using a NanoDrop 2000 spectrophotometer (Thermo Scientific). cDNA was made from 2 μg total RNA with a RevertAid H Minus First Strand cDNA Synthesis Kit (Thermo Scientific) using random hexamer primers. Quantitative real-time PCR was performed using a Roche LightCycler 480 Instrument II (Roche Diagnostics) with iTaq Universal SYBR Green Supermix (Bio-Rad Laboratories) to detect DNA amplification. The following cycling parameters were used: 95°C for 10 min., followed by 40 cycles at 95°C for 15 sec, 60°C for 30 sec and 72°C for 15 sec. The primers used were GapDH_F, GapDH_R, SERF2_F and SERF2_R.

**Western blot**

Wild-type HEK293T cells and SERF2 CRISPR-deletion mutant cells were collected in RIPA buffer (150mM NaCl, 5mM EDTA pH 8.0, 50mM Tris pH 8.0, 1% NP-40, 0.5% sodium deoxycholate and 0.1% SDS) supplemented with Complete Protease Inhibitor Cocktail (25x) (Roche; 11697498001). Cells were kept on ice for 1 hour and sonicated 20 cycles, 30 sec on, 20 sec off. Lysates were spun down at maximum speed at 4°C for 30 min. The protein concentration in the supernatant was determined using a Pierce™ BCA Protein assay Kit (Thermo Fisher Scientific; 23225). 50μg of proteins was separated on a 12% Tris-Glycine acrylamide gel and transferred to a 0.2µm nitrocellulose membrane (Bio-Rad; 1620112). Membranes were incubated in blocking solution (5% BSA or 5% milk in PBS-T 0.1%) for 1 hour. The antibodies used to detect SERF2-HA and mutant SERF2-HA were primary anti-SERF2 antibody (ProteinTech; 11691-1-AP) in 5% BSA at a dilution of 1:1500, or primary anti-HA antibody-ChIP Grade (Abcam; ab9110) at a 1:5000 dilution in 5% milk. PolyQ was detected using primary anti-Polyglutamines antibody (Sigma-Aldrich; Cat# P1874) at a 1:1000 dilution or primary anti-HA antibody-ChIP Grade (Abcam; ab9110) at a 1:5000 dilution in 5% milk. Tubulin was detected using anti-tubulin antibody (Sigma-Aldrich; T6074) at a 1:5000 dilution in 5% milk. Primary antibody incubation was performed overnight at 4°C. Secondary antibodies secondary goat anti-rabbit IgG (H+L)-HRP conjugate antibody (Bio-Rad; 1706515) and secondary goat anti-mouse IgG (H + L)-HRP conjugate antibody (Bio-Rad; 1706516) were used with a 1:10,000 dilution for 1 hour at room temperature. Antibody binding was visualized using Amersham ECL Prime Western Blotting Detection Reagent (GE Healthcare Life Sciences; RPN2236) and imaged using ImageQuant LAS4000 imaging machine (GE Healthcare Life Sciences).

Worms were collected in PBS supplemented with Complete Protease Inhibitor Cocktail (25x) (Roche; 11697498001). Samples were sonicated 15 cycles, 30 sec on, 15 sec off. Lysates were spun down and protein concentrations in the supernatant were determined using a Pierce™ BCA Protein assay Kit (Thermo Fisher Scientific; 23225). 20µg protein with Laemmli buffer (4x) was incubated at 95°C for 10 minutes. Proteins were separated on a 12% Tris-Glycine acrylamide gel and transferred to a 0.2µm nitrocellulose membrane (Bio-Rad; 1620112). Membranes were incubated in blocking solution (5% Milk in PBS-T 0.1%) for 1 hour. MOAG-4 was detected using primary anti-moag-4 antibody (10B1) at a 1:2000 dilution in 5% milk. Primary antibody incubation was performed overnight at 4°C. Secondary antibody incubation and antibody visualization was performed as described for cells.

**HEK cell transfection and immunocytochemistry**

500,000 HEK293T cells/well were seeded on poly-L-Lysine-coated coverslips in 6-wells plates. After 24 hours, cells were transfected using 1mg/mL polyethylenimine (PEI) reagent (Polysciences, 239662) and 1000ng DNA, and grown for 24 hours. Cells were fixed in 4% cold paraformaldehyde for 15 minutes at room temperature and washed twice with PBS. The cells were permeabilized in PBS-Triton X-100 0.1% for 5 minutes at room temperature, washed once with PBS and blocked twice with PBS+ (1x PBS, 0.5% BSA, 0.15% glycine). Coverslips were labeled for 1 hour with primary anti-HA antibody-ChIP Grade (1:1000, Abcam; ab9110) in PBS+ at room temperature. After labeling, coverslips were washed four times with PBS+ and incubated for 1 hour in secondary goat anti-rabbit antibody, Alexa Fluor 546 (1:800, Invitrogen; A11010), in PBS+ at room temperature. Coverslips were washed twice in PBS+ and once in PBS before mounting on glass microscope slides using VECTASHIELD® Antifade Mounting Medium with DAPI (Vector Labs; H-1200). Cells were imaged using a Leica TCS SP8 confocal microscope.

**Filter Trap assay**

The Filter Trap assay was performed as described previously (7). In brief, $2.5\times10^6$ SERF2 CRISPR-deletion mutant cells were co-transfected with 1000ng (mut)HA-74Q DNA and either 3000ng wild-type hSERF2-HA or mutant SERF2-HA DNA using 1mg/mL polyethylenimine (PEI) reagent (Polysciences, 239662). Cells were collected 16 hours after transfection in FTA sample buffer (10 mM Tris-Cl pH 8.0, 150 mM NaCl and 2% SDS) supplemented with Complete Protease Inhibitor Cocktail (25x) (Roche; 11697498001). Fast-prep treatment was performed 5x 20 sec at 4 m/s (MP Biomedicals; 116004500). Total protein was quantified using a Pierce™ BCA Protein Assay Kit (Thermo Fisher Scientific; 23225). The dilution range was 40 µg, 8 µg and 1.6 µg in 100 µl FTA sample buffer supplemented with 1M DTT (1:20) (Sigma; D0632), and incubation was at 95 °C for 5 minutes. A Bio-Dot apparatus (Bio-Rad) was used to trap the aggregates in a 0.2 micron cellulose acetate membrane (Sterlitech; CA023001). Membrane blocking in 5% milk in PBS-T 0.1%, antibody staining and membrane developing were performed as described above for Western blotting. The antibodies used to detect HA-tagged polyQ74 were primary anti-HA antibody-ChIP Grade (1:5000; Abcam; ab9110), or primary anti-polyglutamines antibody (1:1000; Sigma-Aldrich, P1874) in 5% milk in PBS-T 0.1%. Filter Trap results were quantified using ImageJ. In addition, a Western blot was performed with 50µg protein of the cell lysates used for each Filter Trap, to quantify protein expression in the different samples as described above for Western blot. PolyQ74 was detected using primary anti-HA antibody-ChIP Grade (1:5000; Abcam; ab9110) or primary anti-polyglutamines antibody (1:1000; Sigma-Aldrich, P1874) in 5% milk in PBS-T 0.1%. Anti-tubulin (Sigma-Aldrich; T6074) or anti-actin (MP Biomedicals; 08691001) was used as loading control at a 1:5000 dilution.

##### *C. elegans* strains PCR

Standard conditions were used for *C. elegans* culturing at 20ºC (8). Animals were age-synchronized by hypochlorite bleaching and hatched overnight in M9 medium. The L1 animals were plated on nematode growth medium (NGM) plates seeded with OP50 bacteria. The strains that were used are listed in Materials table. The MOAG-4 charge mutant (PHX1173), MOAG-4 control mutant (PHX1099) and MOAG-4 KR mutant (PHX2470) were generated using CRISPR/Cas9 to make the following mutations: PHX1173 K16E, K17E and K23Q, and PHX1099 L18V, A19V and Q21L, and PHX2470 K16R, K17R and K23R (Sunybiotech, China). Mutations were confirmed by PCR and sequencing using the MOAG-4 seq Fw and MOAG-4 seq rev primers.

#### Quantification of aggregates

The numbers of aggregates present in whole worms (L4 stage) were counted using a fluorescence dissection stereomicroscope (Leica, MZ16 FA). For each replicate, aggregates were counted in 20 animals. Statistical analysis was performed using GraphPad Prism 5. One-way ANOVA and Bonferroni tests were used for comparisons. P-values below 0.05 were considered statistically significant. *=$P<0.05$; **=$P<0.01$; ***=$P<0.001$ and ****=$P<0.0001$. All data are represented as mean ± standard deviation.

#### Peptide synthesis and labeling for MST

Peptides were synthesised in-house using an Intavis Multipep RSi synthesizer following Fmoc/tBu solid-phase synthesis strategy. Peptides were subsequently labeled with 5(6)-carboxyfluorescein as follows: 5 equiv. of 5(6)-carboxyfluorescein (0.5M in DMF), 5 equiv. HBTU (0.5M in DMF), 5 equiv. 6-Cl-HOBt (0.5M in DMF) and 10 equiv. NMM (4M in DMF) were added to the resin. After 4h the reaction was stopped and the resin washed. After cleavage from the solid support, the labeled peptides were stored as ether stocks at -20°C. If necessary, crude peptides were purified using RP-HPLC purification protocols to ensure high levels of peptide purity (> 90%).

#### Microscale thermophoresis (MST)

Directly prior to the MST analysis, a peptide aliquot was dried, resolubilized in 1,1,3,3-hexafluoroisopropanol and dried again. The resulting film was dissolved in DMSO, and diluted 1:20 in 10 mM tris Ph 7.6 prior, after which concentration was determined using a NanoDrop (ThermoFisher). Peptides were then diluted to a concentration of 600 nM in 10 mM Tris pH 7.6 with 0.5% Tween-20. SERF2 or mutants thereof were diluted to a concentration of 107 µM in 10 mM Tris, and a 1:1 dilution series produced. This dilution series was then mixed at a 9:1 ratio with 600nm peptide, resulting in a final assay buffer of 10 mM Tris with 0.05% Tween-20 and a final labelled peptide concentration of 60 nM.

Thermophoresis was then performed on a Monolith NT. Automated (NanoTemper, Germany). KD determination was performed using the MO.Affinity Analysis (NanoTemper, Germany) software version 2.3, using a KD model and keeping the target concentration constant at 60 nM.

##### *In vitro* aggregation assay analysis

The ThT kinetic assays were analyzed using the sigmoidal fit function from the 'sicegar' package in R. The fitting algorithm was applied to the average intensity from all replicate experiments. For blank corrections, the minimal measured intensities from all-time series were subtracted from all other intensity values in that series. Subsequently, all ThT amyloid kinetic data were plotted using the 'figureModelCurves' function of the 'sicegar' package in R.

##### Quantitative PCR analysis

All quantitative PCR results represent the mean of n = 2 biological replicates, with error bars representing mean ±SD. Relative transcript levels were quantitated using a standard curve of pooled cDNA solutions. Expression levels were normalized against the endogenous reference gene GapDH.

##### Filter Trap assay analysis

Immunoblots were quantified by densitometry using ImageJ. The relative amount of SDS-insoluble protein was quantified by calculating the ratios (fold change) of the values of wild-type SERF2/SERF2 mutants relative to the empty vector control. All statistical analyses were performed using GraphPad Prism 5. Data were analyzed using one-way ANOVAs, and Bonferroni's tests were used for comparisons. P-values below 0.05 were considered statistically significant. *=P<0.05; **=P< 0.01; ***=P<0.001. All data are represented as mean ± standard deviation.

#### SI Materials table

| REAGENT or RESOURCE | SOURCE | IDENTIFIER |
|---|---|---|
| **Antibodies** | | |
| Rabbit anti-SERF2 antibody | ProteinTech | Cat# 11691-1-AP |
| Goat anti-rabbit IgG (H+L)-HRP Conjugate | Bio-Rad | Cat# 1706515 |
| Mouse anti-tubulin | Sigma-Aldrich | Cat# T6074 |
| Goat Anti-Mouse IgG (H + L)-HRP Conjugate | Bio-Rad | Cat# 1706516 |
| Rabbit anti-HA antibody-ChIP Grade | Abcam | Cat# ab9110 |
| Goat anti-Rabbit IgG (H+L), Alexa Fluor 546 | Invitrogen | Cat# A11010 |
| Mouse anti-actin, monoclonal antibody | MP Biomedicals | Cat# 8691001 |
| Monoclonal Anti-Polyglutamines antibody produced in mouse | Sigma-Aldrich | Cat# P1874 |
| Mouse anti-Moag-4 antibody (10B1) | Lab stocks | N/A |

| | | |
|---|---|---|
| **Chemicals, Peptides, and Recombinant Proteins** | | |
| Isopropyl ß-D-1-thiogalactopyranoside | Promega | Cat# V3951 |
| protease inhibitor cocktail tablets (25x) | Roche | Cat# 11697498001 |
| Benzonase | Merck Millipore | Cat# 70746 |
| Thrombin Protease | GE Healthcare | Cat# 27-0846-01 |
| ATTO 633 NHS-ester 1 mg | ATTO-tec GmbH | Cat# AD633-31 |
| polyethylenimine (PEI) reagent | Polysciences | Cat# 239662 |
| VECTASHIELD® Antifade Mounting Medium with DAPI | Vector Labs | Ca# H-1200 |
| Dithiothreitol | Sigma | Cat# D0632 |
| Trizol | Life Technologies | Cat# 15596-018 |
| recombinant alpha-synuclein | This paper | N/A |
| recombinant amyloid-beta | This paper | N/A |
| recombinant human islet amyloid polypeptide | AnaSpec, Fremont, CA, USA | AS-60804 |
| recombinant wild-type SERF2 | This paper | N/A |
| recombinant SERF2 charge mutant | This paper | N/A |
| recombinant SERF2 KR-mutant | This paper | N/A |

| | | |
|---|---|---|
| **Critical Commercial Assays** | | |
| Q5® Site-Directed Mutagenesis Kit | New England Biolabs Inc. | Cat# E0554S |
| Dulbecco's Modified Eagle Medium; high glucose, pyruvate | Gibco | Cat# 41966052 |
| Bovine Cow Serum | Sigma | Cat# 12133C |
| Penicillin/Streptomycin 10,000 U/mL | Gibco | Cat# 15140122 |
| 0.05% Trypsin-EDTA (1X), Phenol Red | Invitrogen | Cat# 25300-054 |
| Pierce$^{TM}$ BCA Protein Assay Kit | Thermo Fisher Scientific | Cat# 23225 |
| Amersham ECL Prime Western Blotting Detection Reagent | GE Healthcare Life Sciences | Cat# RPN2236 |
| 0.2 uM Cellulose acetate membrane | Sterlitech | Cat# CA023001 |
| SYBR Green Dye | Bio-Rad | Cat# 172-5125 |

| | | |
|---|---|---|
| **Experimental Models: Organisms/Strains** | | |
| *E. coli* competent BL21 (DE3) cells | Lab stocks | N/A |
| NEB 5-alpha Competent *E. coli* | New England Biolabs Inc. | Cat# C2987H |
| HEK293T cells | Lab stocks | N/A |
| SERF2 CRISPR clone 1 HEK293T cells | This paper | N/A |
| SERF2 CRISPR clone 2 HEK293T cells | This paper | N/A |

| **C elegans strains** | | |
|---|---|---|
| N2 | CGC | N2 |
| PHX1099 moag-4(syb1099) | SunyBiotech | PHX1099 |
| PHX1173 moag-4(syb1173) | SunyBiotech | PHX1173 |
| PHX2470 moag-4(syb2470) | SunyBiotech | PHX2470 |
| OW441 moag-4(gk513) I | van Ham et al., 2010 | OW441 |
| OW452 moag-4(tm4909) I | CGC | OW452 |
| Am141 rmIs133 [unc-54p::Q40::YFP]X | Morley et al., 2002 | AM141 |
| OW201 moag-4(gk513) I; rmIs133[P(unc-54)::Q40::YFP] X | van Ham et al., 2010 | OW201 |
| OW202 moag-4(syb1099) I; rmIs133[P(unc-54)::Q40::YFP] X | This paper | OW202 |
| OW203 moag-4(syb1173) I; rmIs133[P(unc-54)::Q40::YFP] X | This paper | OW203 |
| OW204 moag-4(syb1099) I; rmIs126[P(unc-54)::Q0::YFP] V | This paper | OW204 |
| OW205 moag-4(syb1173) I; rmIs126[P(unc-54)::Q0::YFP] V | This paper | OW205 |
| OW453 moag-4(tm4909)I; rIs133[P(unc-54)::Q40::YFP]X | This paper | OW453 |
| OW833 Moag-4 (tm4909)I; rmIs126 [P(unc-54)::Q0::YFP]V | This paper | OW833 |
| OW215 moag-4(syb2470) I; rmIs133[P(unc-54)::Q40::YFP] X | This paper | OW215 |

| **Oligonucleotides** | This paper | N/A |
|---|---|---|
| pGEX primer_F1: CGCTTCTAGAGTCGACGAATTCATGACCCGCGGTAACCAG | This paper | N/A |
| pGEX primer_R1: TTATTCTAGAGCGGCCGCCTACTTGGGTTCCTCCTTC | This paper | N/A |
| SDM primer_F1: GTTCAGGGAAAGCGCCGAGATGACGGGCTTTCTGCTG | This paper | N/A |
| SDM primer_R1: CGAGTCGCTCTGCTCTTCCATATTCTTCTGGCGGGC | This paper | N/A |
| SDM primer_F2: GCTAAGGCAAAGCGCCGAGATGACGGGCTTTCTGCT | This paper | N/A |
| SDM primer_R2: CGCGTCGCTCTGCTTTTTCATATTCTTCTGGCGGGC | This paper | N/A |
| GapDH_F: TCAACGGATTTGGTCGTATTG | This paper | N/A |
| GapDH_R: TCTCGCTCCTGGAAGATGG | This paper | N/A |
| Serf2_F: CCCGCCAGAAGAATATGAAA | This paper | N/A |
| Serf2_R: TCGTTTGCCTTTTTCTGCTT | This paper | N/A |
| Moag-4 seq FW : CCTAAACTACACCTCCTTCC | This paper | N/A |
| Moag-4 seq rev : GAGAAGGAGACAGAGGCATT | This paper | N/A |
| Serf2_CRISPR_F: CACCGTATGAAAAGCAGAGCGACT | This paper | N/A |
| Serf2_CRISPR_R: AAACAGTCGCTCTGCTTTTTCATAC | This paper | N/A |

| **Recombinant DNA** | | |
|---|---|---|
| pGEX-4t-1 vector | Sigma | GE28-9545-49 |
| pENG958 [GST::SERF2] | This paper | N/A |
| pENG929 [GST::SERF2 charge mutant] | This paper | N/A |
| pENG934 [empty vector::HA] | This paper | N/A |
| pENG743 [SERF2::HA] | This paper | N/A |
| pENG902 [SERF2 charge mutant::HA] | This paper | N/A |
| pENG924 [SERF2 control mutant::HA] | This paper | N/A |
| pSpCas9(BB)-2A-GFP (PX458) vector | Addgene | Cat# 48138 |
| pENG748 [HA74Q] | gift from prof. Dr. H.H. Kampinga | N/A |
| muHA74Q | This paper | N/A |

| Software and Algorithms | | |
|---|---|---|
| Image Studio Lite Version 5.2 sofware | Open source | N/A |
| HS Pro control manager software | Tecan | N/A |
| Array-Pro® Analyzer software | Tecan | N/A |
| ImageJ | Open source | N/A |
| GraphPad Prism 5 | Graphpad | N/A |

| Other | | |
|---|---|---|
| GSTrap 4B column | GE Healthcare | Cat# 28-4017-45 |
| HiTrap Benzamidine FF column | GE Healthcare | Cat# 17-5143-02 |
| Sephadex G-25 column | GE Healthcare | Cat# 17-0851-01 |
| HS 400™ Pro Hybridization Station | Tecan | N/A |
| Powerscanner Microarray | Tecan | N/A |
| ImageQuant LAS4000 imaging machine | GE Healthcare Life Sciences | N/A |
| FastPrep 24 Instrument | MP Biomedicals | Cat# 116004500 |
| 48-well Bio-Dot microfiltration system | Bio-Rad | Cat# 1703938 |
| Leica TCS SP8 confocal microscope | Leica | N/A |
| Grasshopper3 4.1 MP Mono USB3 Vision camera | Point Grey | GS3-U3-41C6M-C |
| 16mm High resolution lens | Edmund Optics | Cat # 86-571 |
| Tensor 27 FTIR spectrometer | Bruker | N/A |

### SI Figures S1-S5

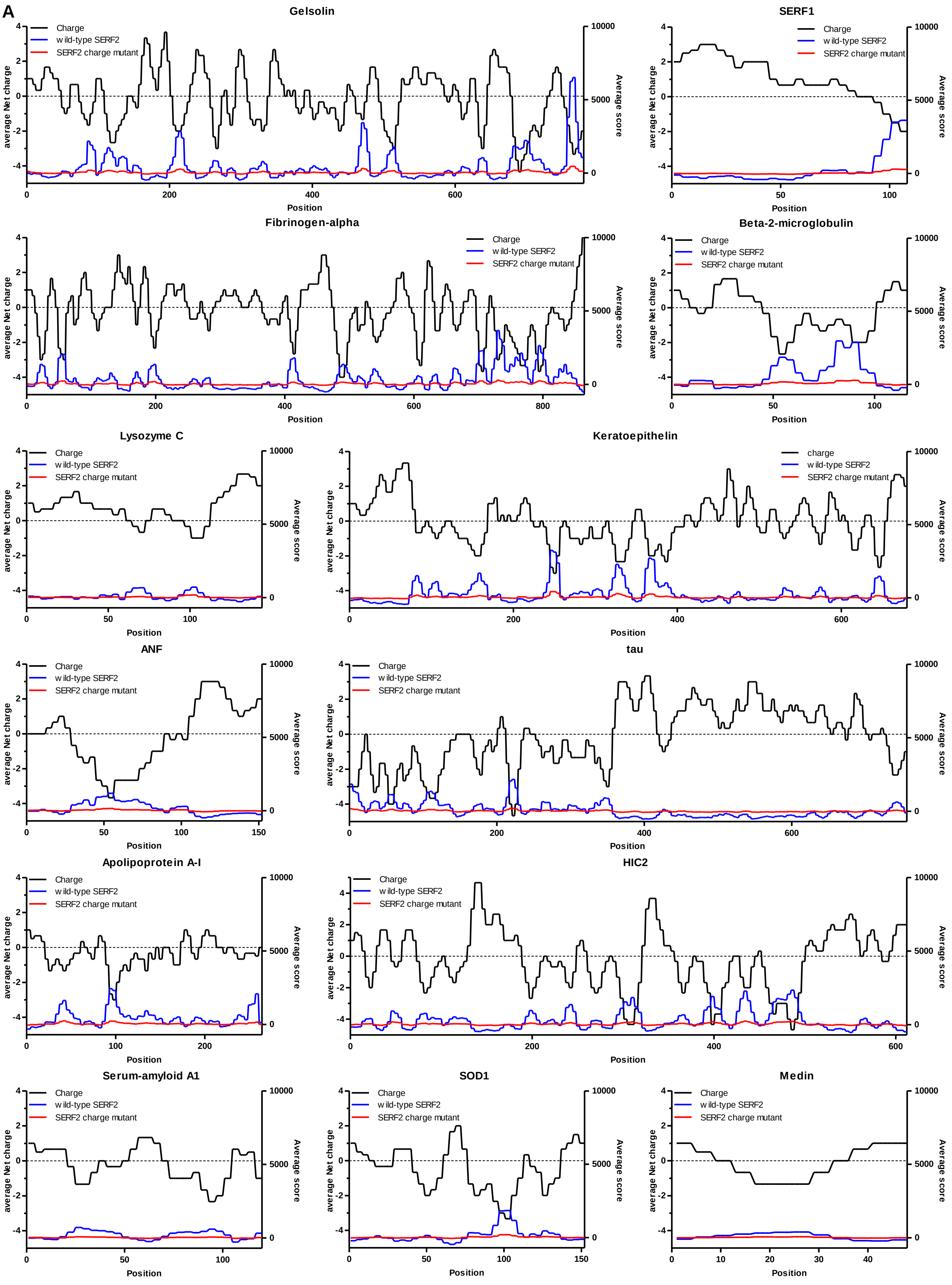

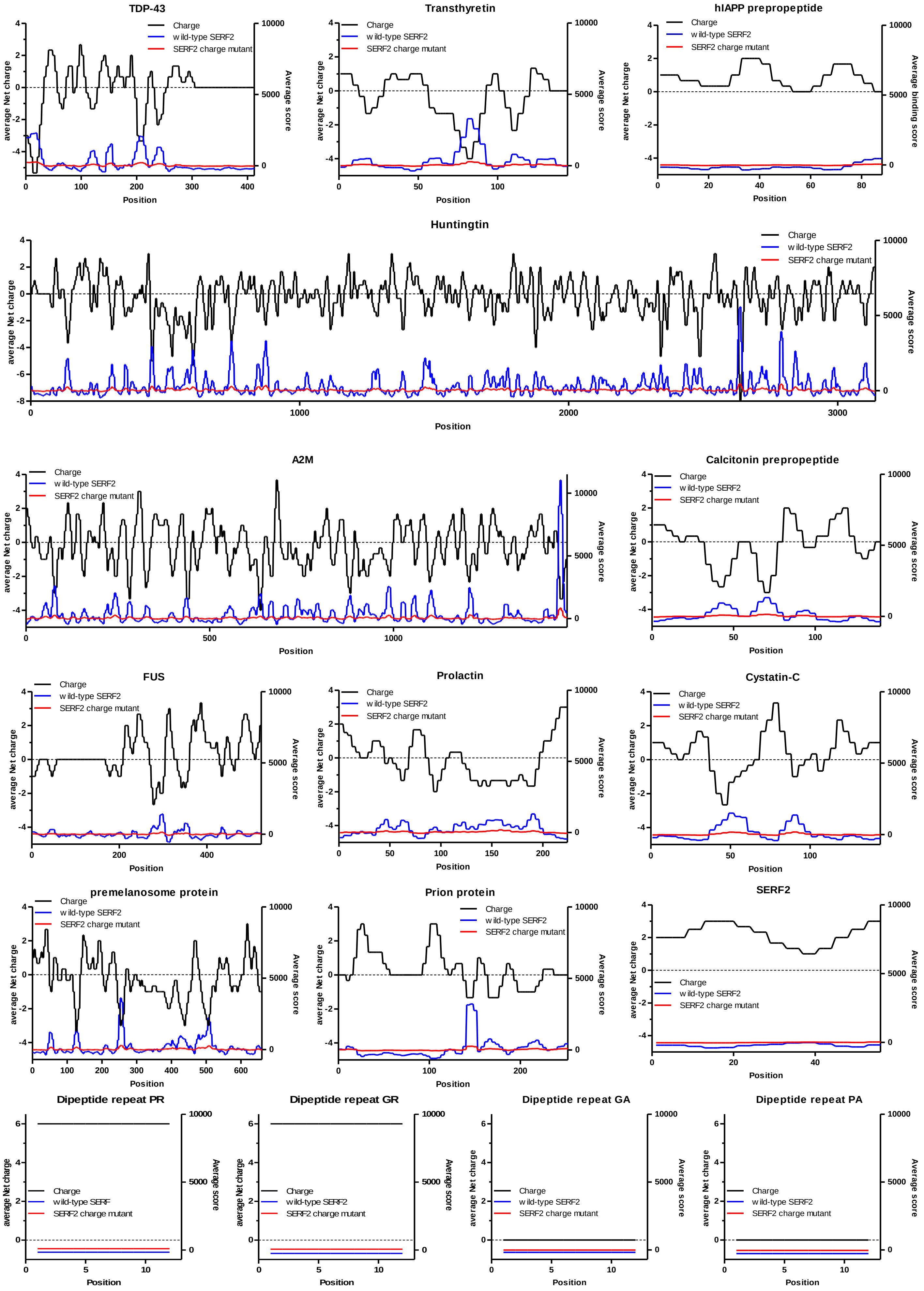
TDP-43
Transthyretin
hIAPP prepropeptide
Huntingtin
A2M
Calcitonin prepropeptide
FUS
Prolactin
Cystatin-C
premelanosome protein
Prion protein
SERF2
Dipeptide repeat PR
Dipeptide repeat GR
Dipeptide repeat GA
Dipeptide repeat PA
Charge
wild-type SERF2
SERF2 charge mutant
wild-type SERF
average Net charge
Average score
Average binding score
Position

**B**

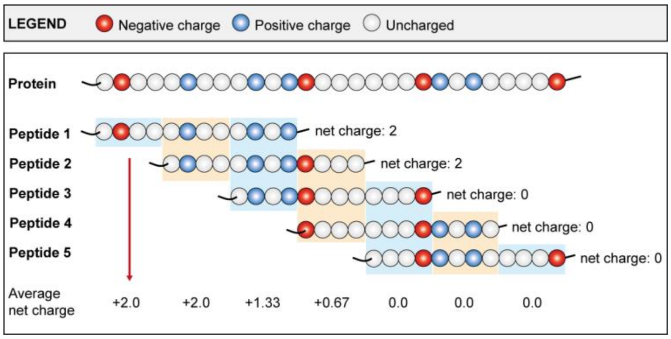


**Supplemental Figure S1.** (A) Charge (black lines) distribution of all full-length proteins present on the peptide micro-array, together with the wild-type (blue lines) and mutant (red lines) average binding score. (B) Schematic representation of the average net charge calculation per residue, corresponding to Figure 2 and Dataset S5.

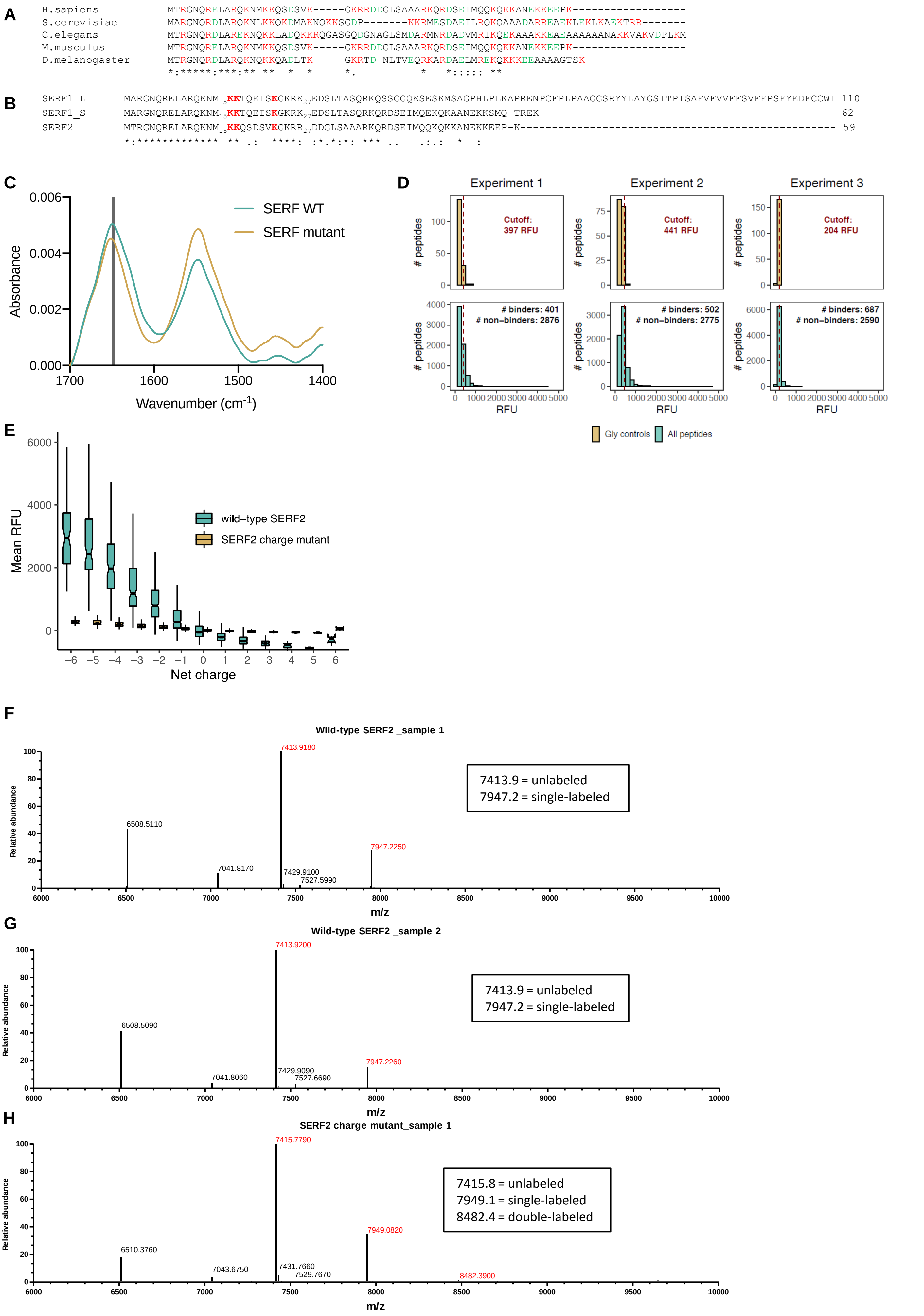

A
H.sapiens MTRGNQRELARQKNMKKQSDSVK-----GKRRDDGLSAAARKQRDSEIMQQKQKKANEKKEEPK-----------------
S.cerevisiae MARGNQRDLARQKNLKKQKDMAKNQKKSGDP-------KKRMESDAEILRQKQAAADARREAEKLEKLKAEKTRR-------
C.elegans MTRGNQRDLAREKNQKKLADQKKRQGASGQDGNAGLSMDARMNRDADVMRIKQEKAAAKKEAEAAAAAAANAKKVAKVDPLKM
M.musculus MTRGNQRELARQKNMKKQSDSVK-----GKRRDDGLSAAARKQRDSEIMQQKQKKANEKKEEPK-----------------
D.melanogaster MTRGNQRDLARQKNQKKQADLTK-----GKRTD-NLTVEQRKARDAELMREKQKKKEEAAAAGTSK----------------
B
SERF1_L MARGNQRELARQKNM15KKTQEISKGKRK27EDSLTASQRKQSSGGQKSESKMSAGPHLPLKAPRENPCFPLPAAGGSRYYLAYGSITPISAFVFVVFFSVFFPSFYEDFCCWI 110
SERF1_S MARGNQRELARQKNM15KKTQEISKGKRK27EDSLTASQRKQRDSEIMQEKQKAANEKKSMQ-TREK--------------------------------------------- 62
SERF2 MTRGNQRELARQKNM15KKQSDSVKGKRR27DDGLSAAARKQRDSEIMQQKQKKANEKKEEP-K------------------------------------------------ 59
C
Absorbance
Wavenumber (cm-1)
SERF WT
SERF mutant
D
Experiment 1
Experiment 2
Experiment 3
Cutoff: 397 RFU
Cutoff: 441 RFU
Cutoff: 204 RFU
### binders: 401
### non-binders: 2876
### binders: 502
### non-binders: 2775
### binders: 687
### non-binders: 2590
### peptides
RFU
Gly controls
All peptides
E
Mean RFU
Net charge
wild-type SERF2
SERF2 charge mutant
F
Wild-type SERF2 _sample 1
7413.9 = unlabeled
7947.2 = single-labeled
6508.5110
7041.8170
7413.9180
7429.9100
7527.5990
7947.2250
Relative abundance
m/z
G
Wild-type SERF2 _sample 2
7413.9 = unlabeled
7947.2 = single-labeled
6508.5090
7041.8060
7413.9200
7429.9090
7527.6690
7947.2260
Relative abundance
m/z
H
SERF2 charge mutant_sample 1
7415.8 = unlabeled
7949.1 = single-labeled
8482.4 = double-labeled
6510.3760
7043.6750
7415.7790
7431.7660
7529.7670
7949.0820
8482.3900
Relative abundance
m/z

I

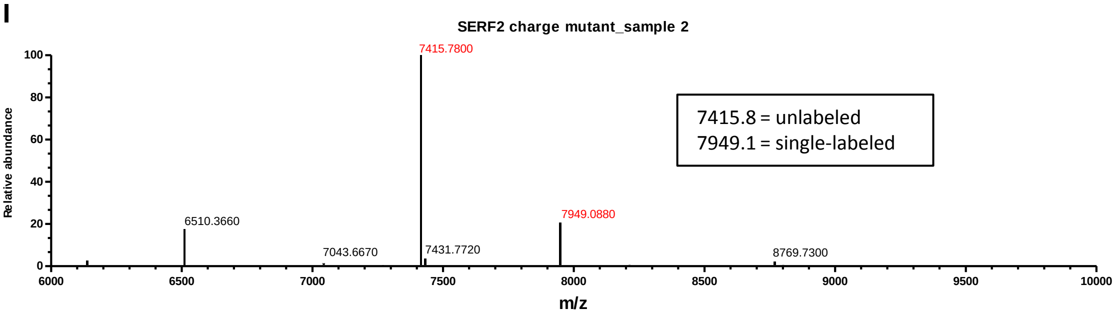


**Supplemental Figure S2.** (A) Alignment of human SERF2 protein with orthologs in yeast *Saccharomyces cerevisiae*, *Caenorhabditis elegans*, *Mus Musculus* and fruit fly *Drosophila melanogaster*, showing the conservation of the N-terminal region. (B) Amino acid sequences of three human SERF proteins, showing the conservation of the charged residues in the N-terminal region. (C) Secundary structure determination of wild-type SERF2 and SERF2 charge mutant proteins. A wavenumber of 1646-1650 $cm^{-1}$ (gray bar) corresponds to random coil. (D) Histograms of SERF2 charge mutant binding to non-control peptides (top panel) and Glycine control peptides (bottom panel). Dashed line indicates the cutoff used for distinguishing binders from non-binders. (E) Boxplots showing binding of SERF2 charge mutant and wild-type versus peptide net charge. The upper whiskers indicate the largest value no further than 1.5 times the Inter-Quartile Range (IQR) from the upper hinge, the lower whiskers show the smallest value at least 1.5 * IQR from the lower hinge. Notches extend 1.58 * IQR / sqrt(n) from either side of the median and represent a 95% confidence interval for the median value. (F-I) Mass Spectrometry analysis of purified, ATTO633-labelled wtSERF2 and SERF2 charge mutant.

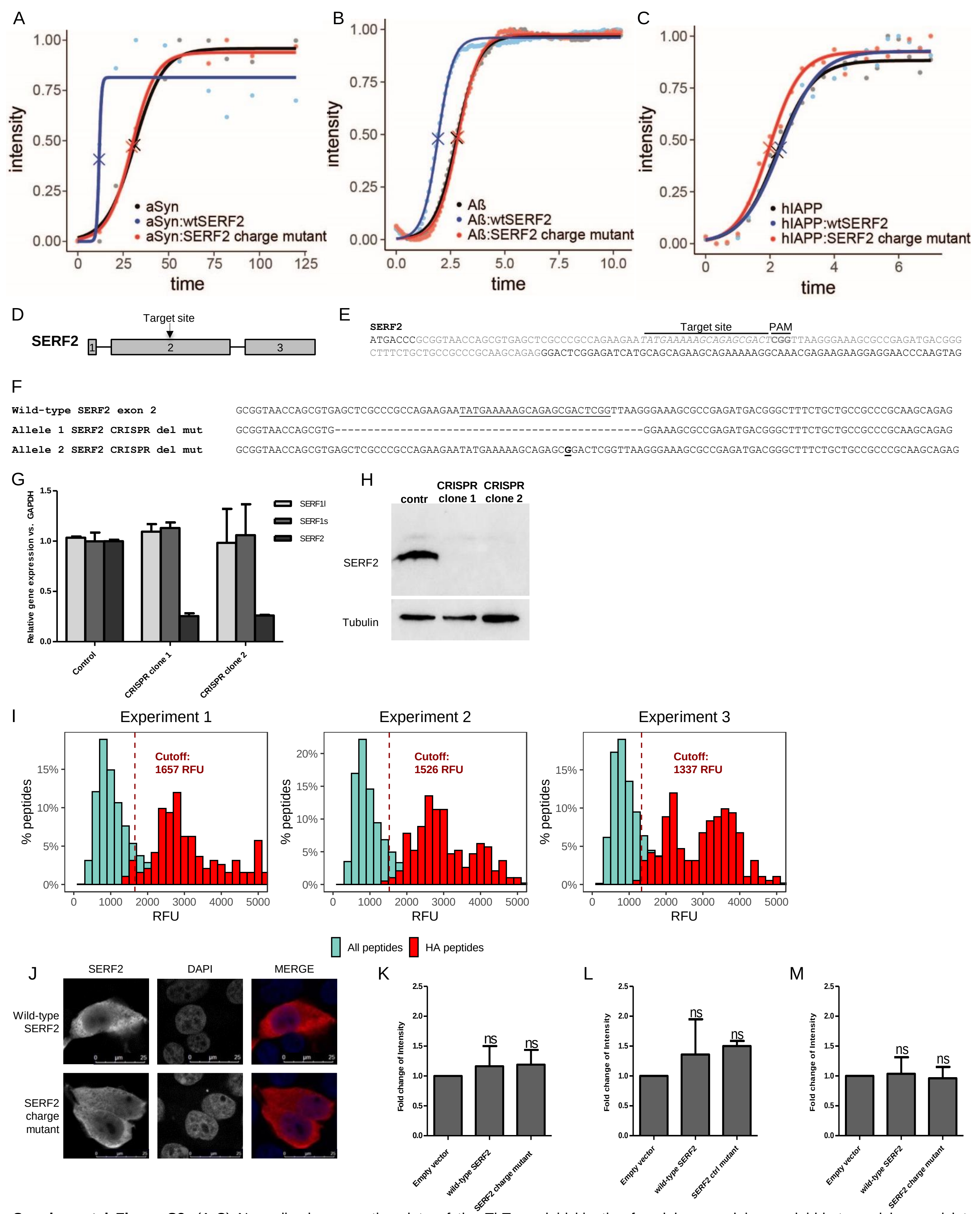


**Supplemental Figure S3.** (A-C) Normalized aggregation data of the ThT amyloid kinetics for alpha-synuclein, amyloid-beta and human islet amyloid polypeptide in the presence of either wild-type SERF2 or mutant SERF2, corresponding to figures 5C, 5D and 5F. (D+E) Generation of *SERF2* CRISPR-deletion mutant cell lines; targeting strategy to generate mutations in the second exon of *Serf2*. (F) Sequencing results for the *SERF2* CRISPR-deletion mutant cell line. (G) qPCR analysis of the *SERF2* CRISPR clone 1 and 2, showing specific loss of *Serf2* gene expression in both *SERF2* mutant cell lines. N=2 per cell line. qPCR primers were designed outside the target area. Data are presented as the average SD-fold reduction in samples analysed in triplicate. Only CRISPR clone 2 was used in this paper. (H) Western blot analysis of SERF2 CRISPR-deletion mutant cell lines. Only CRISPR clone 2 was used in this paper. (I) Density plot of wild-type SERF2 binding intensities to HA peptides versus non-control peptides. Dashed line indicates the cutoff used for distinguishing binders from non-binders. Histograms marked in red represent HA-peptides. (J) Localization of wild-type SERF2 and SERF2 charge mutant protein in the *SERF2* mutant CRISPR cell line. (K-M) PolyQ expression corrected for tubulin expression corresponding to figures 5I-K. Data are represented as mean ± SD and significance was calculated using a one-way ANOVA followed by a post-hoc Bonferroni multiple comparisons test (K) p=0.7048, (L) p=0.1747, (M) p=0.9283.

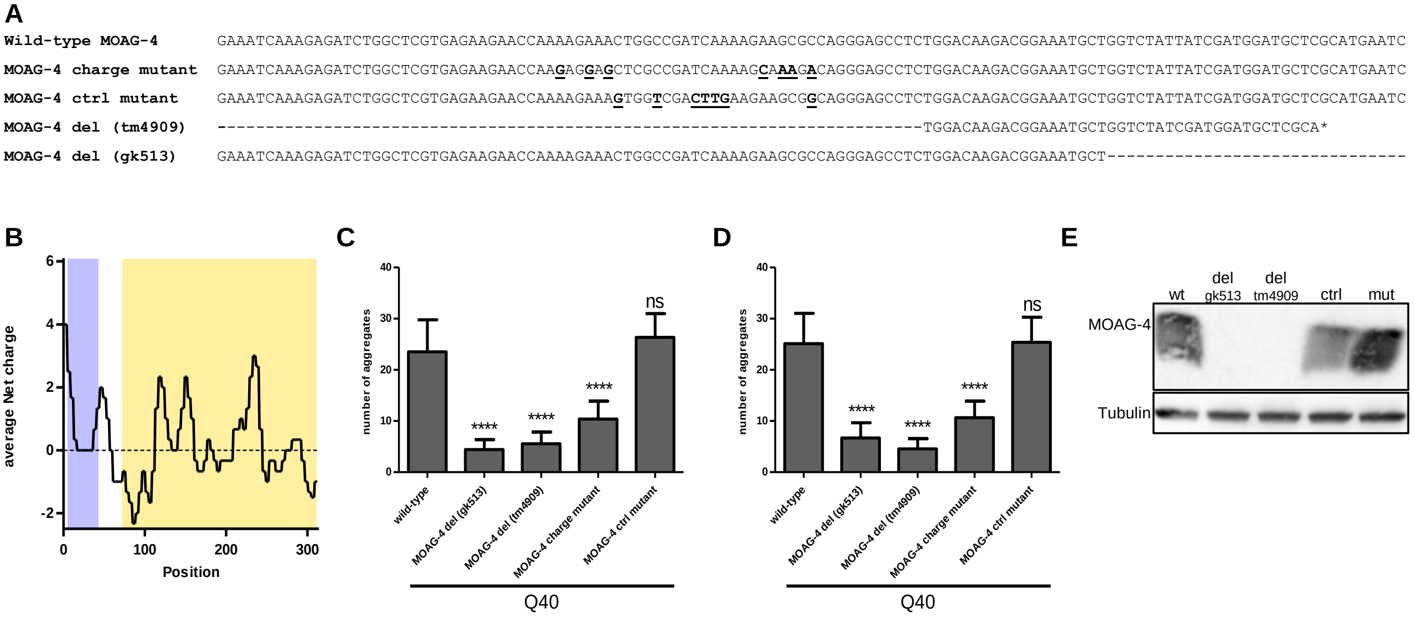


**Supplemental Figure S4.** (A) Sequencing results for MOAG-4 wild-type, mutant and deletion strains. (B) Charge distribution of the Q40 (blue)::linker (white)::YFP (yellow) protein expressed in *C. elegans* (C+D) Replicate quantifications of the number of aggregates in Q40 worms with expression of either wild-type *moag-4*, *moag-4* charge mutant, *moag-4* ctrl mutant or *moag-4* deletion. The results shown are representative experiments of three biological replicates of n=20 worms in L4 stage. Data are represented as mean ± SD and significance was calculated using a one-way ANOVA followed by a post-hoc Bonferroni multiple comparisons test. (C) replicate 2 of figure 6C, p<0.0001, (D) replicate 3 of figure 6C, p<0.0001. ****p < 0.0001. (E) Western blot of MOAG-4 protein levels in N2 wild-type (wt) worms, MOAG-4 deletion (gk513 and tm4909), MOAG-4 control mutant (ctrl) and MOAG-4 charge mutant (mut) strains.

**A**

**Wild-type MOAG-4** GAAATCAAAGAGATCTGGCTCGTGAGAAGAACCAAAAGAAACTGGCCGATCAAAAGAAGCGCCAGGGAGCCTCTGGACAAGACGGAAATGCT

**MOAG-4 KR-mutant** GAAATCAAAGAGATCTGGCTCGTGAGAAGAACCAAA**G**GA**G**ACTGGCCGATCAAAAG**CGT**CG**A**CAGGGAGCCTCTGGACAAGACGGAAATGCT

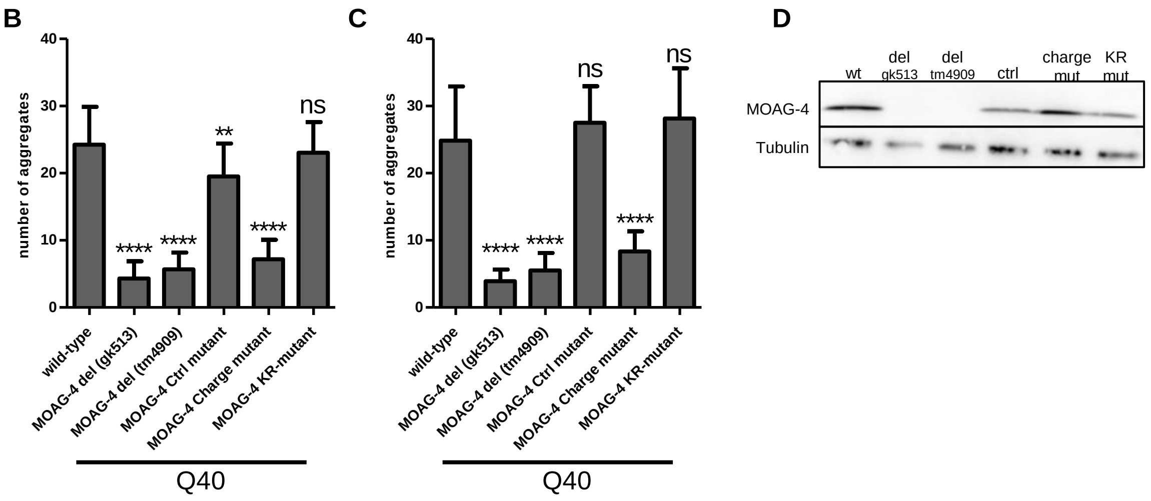


**Supplemental Figure S5.** (A) Sequencing results for MOAG-4 wild-type and KR-mutant strains. (B+C) Replicate quantifications of the number of aggregates in Q40 worms with expression of either wild-type *moag-4*, *moag-4* charge mutant, *moag-4* ctrl mutant , *moag-4* KR-mutant or *moag-4* deletion. The results shown are representative experiments of three biological replicates of n=20 worms in L4 stage. Data are represented as mean ± SD and significance was calculated using a one-way ANOVA followed by a post-hoc Bonferroni multiple comparisons test. (B) replicate 2 of figure 7F, $p<0.0001$, (C) replicate 3 of figure 7F, $p<0.0001$. **p < 0.01; ****p < 0.0001. (D) Western blot of MOAG-4 protein levels in N2 wild-type (wt) worms, MOAG-4 deletion (gk513 and tm4909), MOAG-4 control mutant (ctrl), MOAG-4 charge mutant (charge mut) and MOAG-4 KR-mutant (KR mut) strains.
